# A Two-Arm Metabolic-Efflux Adaptation Framework in *Klebsiella pneumoniae* under Mixed Pharmaceutical Exposure: *rutA*-Linked Oxidative Entry and *rutR*-Associated Regulation

**DOI:** 10.64898/2026.07.11.738005

**Authors:** Shilpa Sinha, Partha Barman, Debopriya Haldar, Ranadhir Chakraborty

## Abstract

Chemically complex pharmaceutical mixtures in wastewater and sludge can affect microbial adaptation; however, the responses to different co-occurring compounds have not been elucidated well. In this study, the adaptation of a strain derived from hospital sludge, *Klebsiella pneumoniae* SS02, to 17α-ethinylestradiol (EE2), warfarin sodium, and their combination has been studied. The organism grows under all three conditions, and pre-exposure experiments show induction and cross-induction to substrates. UHPLC MS/MS analyses demonstrated that there is conditional depletion of the parent compound EE2 by ∼15% at 36 h post-treatment compared to initial concentrations, but not for the abiotic and non-adapted controls. The rate of warfarin sodium depletion was approximately ∼30% within 36 h and was in accordance with first order kinetics (k = 0.0102 /h; t□/□= 67.9 h). Under the combined treatment regime, there was a delay in warfarin sodium depletion, suggesting staged substrate consumption. Growth inhibition with efflux inhibitors confirmed transport-driven tolerance. A genome-based study revealed the coordinated response strategy that involved a proposed flavin-dependent monooxygenase (RutA), an oxidative entry into the pathway; redox processing linked to Hpa; aromatic metabolism through β-ketoadipate pathway; and RND efflux system. The structural study additionally supported ligand-mediated decrease in DNA binding affinity of RutR, which is in agreement with de-repression of the substrate-activated regulatory mechanism. All these findings lead to the development of a dual-strategy for adaptation model in which oxidative modification and efflux-mediated protection work together under the influence of a mixture of pharmaceuticals.

## 1. Introduction

The occurrence of pharmaceuticals in wastewater, sludge, and the receiving environment as mixture systems rather than as pure substances has been increasing (Patel et al., 2019; Santos et al., 2010; Tang et al., 2021a). In these mixture systems, the endocrine disrupting compound 17α-ethinylestradiol (EE2) and the therapeutic drug warfarin sodium are examples of two structurally different groups of compounds that can be present together in the waste streams from hospitals and municipalities (Tang et al., 2021a, 2021b). Although the transformation processes for certain pharmaceuticals have been identified, not enough research has been conducted regarding the response of microorganisms exposed to mixed pharmaceuticals (Joss et al., 2006; Patel et al., 2019).

Hospitals’ sludge can be regarded as a chemically diverse niche which is highly enriched with various medicines, metabolites, and stressors, thus allowing for the selection of microbes which can metabolically adapt to such conditions. It has been previously demonstrated that biotransformation of steroidal estrogens typically follows oxidative activation, after which the compound is further incorporated into core metabolism (Chen et al., 2018, 2017; Hashem et al., 2025; Hsiao et al., 2021; Ibero et al., 2020; Tian et al., 2022), while biotransformation pathways of anticoagulants such as warfarin sodium have been poorly characterized (Fernández et al., 2014; Gómez-Canela et al., 2014). Furthermore, the degree to which both compounds overlap in their metabolic and regulatory responses is hardly understood.

For this study, a bacterium isolated from hospital sludge was characterized, namely *Klebsiella pneumoniae* SS02, grown with EE2, and tested for sensitivity to EE2, warfarin sodium, and a mixture of these two compounds. Growth, substrate consumption and potential response strategies were studied using physiological tests, analytical quantification, genome-based studies, and structural modeling. The study suggests an adaptive strategy based on the use of a common framework of oxidative transformation of xenobiotics, metabolism of aromatic compounds, regulatory changes, and efflux-mediated resistance. The integration of phenotypic, analytical, genomic and structural approaches allowed obtaining insights into the response of environmental bacteria to complex mixtures of chemically diverse pharmaceuticals in sludge environments.

## 2. Materials and Methods

### 2.1. Chemicals and media

17α-ethinylestradiol (EE2) (E4876-100MG) was purchased from Sigma-Aldrich, India. Stock solution of EE2 was prepared in ethanol (50 mg mL^-1^) and stored at 4 °C protected from light. Warfarin sodium (C_19_H_15_NaO_4_, MW=330.31, >98.0% purity) was obtained from TCI Chemicals, India. A 10 mg mL^-1^ stock solution of warfarin sodium was prepared in distilled water and stored at 4 °C.

Luria broth (LB) (HiMedia M575-500G) was employed for routine culturing and inoculum preparation. Minimal Salt Medium (MSM) was used for enrichment, and growth studies (Aneja, 2018). For preparing agar plates, MSM or LB was supplemented with agar (15 g L^-1^).

### 2.2. Sample collection, enrichment, and isolation of strain SS02

Sludge was collected aseptically from North Bengal Medical College and Hospital (NBMCH), Siliguri, West Bengal, India, and processed immediately for enrichment. Enrichment was performed in 100 mL MSM, with EE2 (10 mg L^-1^) as the sole carbon source and incubated at 30 °C at 100 rpm for 7-8 days. Sub-culturing was performed into fresh EE2-MSM under identical conditions. After final enrichment, cultures were serially diluted and plated onto EE2-MSM agar. Morphologically distinct colonies were purified by repeated streaking, and one purified isolate showing reproducible growth on EE2-MSM was selected and designated SS02. Enrichment media composition is given in Table S1.

The ability of the isolate SS02 to utilize a structurally distinct pharmaceutical substrate, a purified colony was streaked on MSM agar containing warfarin sodium (20 mg L^-1^) as sole carbon source. Growth was further confirmed in liquid MSM with warfarin sodium (20 mg L^-1^). Abiotic controls and heat-killed controls were included to distinguish biotic transformation-associated loss and non-biological disappearance. Composition of warfarin sodium-MSM agar is given in Table S2.

### 2.3. Phenotypic assessment of EE2 and warfarin sodium response in strain SS02

Freshly grown SS02 culture was inoculated in MSM containing glucose and incubated at 30 °C till OD_600_ ≈ 0.6. Cells were harvested, washed with 0.85% NaCl, and used as 1% inoculum for MSM-based assays. Composition of growth media is given in Table S3.

#### 2.3.1. Growth assay with EE2 as sole carbon source

MSM was supplemented with EE2 (10 mg L^-1^) as sole carbon source, inoculated with washed cells, and distributed equally into sterile Erlenmeyer flasks. All the flasks were incubated at 30 °C at 100 rpm for 60 h. Samples were taken at 0, 12, 24, 36, 48, and 60 h.

#### 2.3.2. Growth assay with warfarin sodium as sole carbon source

MSM supplemented with warfarin sodium (20 mg L^-1^) as sole carbon source was inoculated and distributed similarly. The flasks were incubated at 30 °C at 100 rpm for 60 h. Samples were collected at 0, 12, 24, 36, 48, and 60 h.

#### 2.3.3. Growth assay with EE2 + warfarin sodium as combined carbon sources

To examine growth under mixed pharmaceutical exposure, MSM was supplemented with EE2 (10 mg L^-1^) and warfarin sodium (20 mg L^-1^) together and inoculated as above. The inoculated media were incubated at 30 °C at 100 rpm for 60 h.

Growth was expressed as log CFU mL^-1^.

#### 2.3.4. Induction of EE2 growth response by EE2 and warfarin sodium pre-exposure

To assess whether prior exposure altered EE2-supported growth, SS02 cells were separately pre-exposed to EE2 at 0.5, 1.0, and 2.5 mg L^-1^, while warfarin sodium at 1, 2, and 5 mg L^-1^. A non-induced culture was maintained as control. After pre-exposure, cells were transferred to fresh EE2-MSM and incubated at 30 °C, 100 rpm, for 12 h. Growth was expressed as log CFU mL^-1^ and compared with the non-induced control.

#### 2.3.5. UV-Vis monitoring of residual warfarin sodium in single and combined-substrate medium

A calibration curve for warfarin sodium was prepared by measuring absorbance at 309 nm for standards ranging from 0.0025 to 0.025 mg mL^-1^ in MSM using a UV-Vis Spectrophotometer (SPECTROstar^Nano^, BMG LABTECH). Residual warfarin sodium was monitored in MSM containing warfarin sodium, 20 mg L^-1^, alone or combined with EE2, 10 mg L^-1^. Cultures were incubated at 30 °C, 100 rpm, and at 12 h intervals, cells were removed by centrifugation. Diluted supernatants were analyzed at 309 nm (Babhair et al., 1985). Residual concentrations were calculated from the regression equation.

Depletion kinetics were fitted to a first-order model by plotting ln (C) against time, where C is the concentration. The apparent first order rate constant (k) was obtained from the slope and half-life (t_1/2_) was calculated as 0.693 k^-1^. Model fit was evaluated using the regression R^2^ and residual inspection (Alexander, 1999).

In combined EE2 + warfarin sodium medium, SS02 growth and residual warfarin sodium were monitored in parallel. This allowed comparison between cell count increase and warfarin sodium depletion under mixed pharmaceutical exposure. Because UV-analysis was used only for warfarin sodium, targeted UHPLC-MS/MS analysis was performed separately to quantify parent EE2 in selected culture supernatants.

#### 2.3.6. Targeted UHPLC-MS/MS analysis of EE2

Targeted UHPLC-MS/MS analysis was performed in multiple reaction monitoring (MRM) mode to quantify parent EE2 in selected culture supernatants using an Acquity reverse-phase UHPLC system coupled to a Waters Xevo tandem quadrupole detector with an electrospray ionization source. Separation was performed on a Waters BEH C18 column, 50 x 2.1 mm, 1.7 µm, using acetonitrile and aqueous buffer containing 5% methanol and 0.2% formic acid as mobile phases. The injection volume was 2 µL and the flow rate was 0.350 mL min^-1^. EE2 was monitored using the EE2 pure compound MRM signal at approximately 0.91 min and quantified by external calibration with 1/x-weighted linear regression based on peak area response. EE2 concentrations in SS02 culture supernatants were calculated from the processed calibration curve. To visualize parent EE2 depletion, supernatants collected at 9 and 36 h were compared with 0 h EE2 reference sample, and reference-to-sample concentration change was plotted. Abiotic and *E. coli* K12 (MTCC1302) controls were included for comparison.

#### 2.3.7. PA***β***N-based efflux inhibition assay under elevated warfarin sodium stress

An efflux inhibition assay was performed to assess whether efflux activity contributes to warfarin sodium tolerance in strain SS02. Since SS02 showed growth in warfarin sodium supplemented MSM, the assay was conducted under both normal and elevated warfarin sodium stress.

Strain SS02 was cultured in MSM supplemented with warfarin sodium at 20 mg L^-1^ and 200 mg L^-1^ in the presence and absence of PAβN, a known efflux pump inhibitor that targets RND-type transporters. The working concentration of PAβN was 10 µg mL^-1^ (Chatterjee et al., 2024). Cultures were incubated at 30 °C at 100 rpm. Samples were collected at 0, 6, 12, 18, and 24 h, serially diluted in sterile 0.85% NaCl, plated on Luria Agar (LA), and expressed as log CFU mL^-1^.

### 2.4. Whole-genome sequencing, annotation, and phylogenomic placement of SS02

Strain SS02 was deposited at Microbial Type Culture Collection (MTCC), Chandigarh under accession number MTCC13772. Whole-genome sequencing was performed using the Illumina HiSeq platform. Genome annotation was carried out using NCBI Prokaryotic Genome Annotation Pipeline (PGAP) and Rapid Annotation using Subsystem Technology (RAST) (Aziz et al., 2008; Bergey, 1994; Tatusova et al., 2016). Whole-genome-based phylogeny was inferred using Type Strain Genome Server (TYGS), and independently reproduced in KBase. Concordant TYGS and KBase tree topology was used to confirm the taxonomic position of strain SS02 (Arkin et al., 2016; Meier-Kolthoff and Göker, 2019).

### 2.5. Genome mining for EE2, warfarin sodium, aromatic metabolism, and efflux-related genes

Whole genome annotation was performed using RAST and eggNOG-mapper v2 to identify genes potentially linked to EE2 and warfarin sodium response (Aziz et al., 2008; Cantalapiedra et al., 2021). Candidate genes were screened for monooxygenases, oxidoreductases, dehydrogenases, hydrolases, ring-cleavage enzymes, redox partners, aromatic-ring processing, central carbon-funneling routes, and efflux-related determinants.

### 2.6. Conserved domain analysis of RutA

The RutA protein sequence of strain SS02 was retrieved from the annotated genome. Conserved domain analysis was performed to verify the functional architecture of RutA and to assess its relevance as a putative oxidative entry-point enzyme for EE2 and warfarin sodium response. The RutA sequence was analysed using NCBI Conserved Domain Search, and eggNOG-mapper v2 (Cantalapiedra et al., 2021; Wang et al., 2023). The analysis mainly focussed on identifying conserved domains associated with flavin-dependent monooxygenase activity, FMN/FAD binding, pyrimidine monooxygenase-like function, and oxidative ring activation. Domain hits were evaluated based on domain identity, accession number, E-value, conserved catalytic features, and predicted functional annotation.

### 2.7. STRING protein-protein interaction (PPI) network analysis

A PPI network was constructed in STRING v11 with *Klebsiella* as the reference background organism to examine functional connectivity among genes implicated in warfarin sodium and EE2 metabolism and tolerance in SS02 (Szklarczyk et al., 2019). Candidate proteins Rut-associated proteins, lower-pathway and auxiliary metabolic proteins, and efflux transporters. Proteins were mapped using locus tags or protein IDs, and interactions were inferred using STRING evidence channels, including curated databases, experiments, gene neighborhood, gene co-occurrence, and co-expression. Only high-confidence interactions were retained. The network was clustered into five modules using k-means and exported for visualization (Szklarczyk et al., 2019).

### 2.8. Construction of the hypothesized EE2-warfarin sodium metabolic and regulatory pathway

A hypothesized EE2-warfarin sodium response pathway was constructed by integrating growth assays, induction response, warfarin sodium depletion data, PAβN-based efflux inhibition, genome mining, conserved domain analysis, STRING network analysis, and structural modelling. Candidate genes were grouped into oxidative entry, aromatic/ring-processing, central carbon-funneling, efflux tolerance, and regulatory modules.

### 2.9. RutR-operator DNA docking for OFF and ON regulatory-state modelling

To evaluate whether EE2 and warfarin sodium binding could alter RutR-DNA interaction and provide a regulatory explanation for the induction response observed in strain SS02, RutR-operator DNA docking was performed. RutR is a transcriptional regulator associated with the RutA-linked oxidative response (Nguyen Le Minh et al., 2015). The RutR protein model was docked with the predicted operator DNA sequence to generate the apo RutR-DNA complex, considered as the OFF-state model. For ON-state modelling, EE2 or warfarin sodium-bound RutR was docked with the operator DNA using HADDOCK (van Dijk, 2006) to assess whether ligand binding reduced RutR-DNA engagement. The resulting clusters were compared based on binding energy, buried surface area, overall complex geometry, and protein-DNA interface features. Protein-DNA interactions were analyzed using PLIP and PyMOL. Hydrogen bonds, hydrophobic interactions, salt bridges, and the number of protein atoms within 4 Å of DNA were recorded for each model.

#### 2.9.1. In silico mutation of protein-DNA contact residues

To further assess the contribution of selected protein-DNA contact residues, residues identified from PLIP analysis were subjected to in silico mutation using PyMOL. Contact residues were selected based on their involvement in hydrogen bonds, hydrophobic interactions, or salt bridges with the operator DNA, with particular attention to residues forming phosphate-backbone interactions. Each selected residue was mutated individually in the RutR-DNA model using the PyMOL mutagenesis tool. The most favourable rotamer was selected based on local geometry and absence of severe steric clashes. Mutated structures were saved separately and compared with the corresponding wild-type model.

This mutation analysis was used as a structural perturbation approach to support the role of selected contact residues in RutR-DNA binding, and not as a substitute for experimental mutagenesis.

### 2.10. Molecular docking of RutA and efflux proteins with EE2 and warfarin sodium

Molecular docking was performed to assess the binding compatibility of RutA with EE2 and warfarin sodium, and of efflux-associated transporters, including MdtA, MdtB, and MdtC, with warfarin sodium. Protein structures were retrieved from the AlphaFold Protein Structure Database and prepared using AutoDock Tools by removing water molecules, adding polar hydrogens, assigning charges, and converting to PDBQT format (Morris et al., 2008; Varadi et al., 2022). EE2 and warfarin sodium structures were obtained from PubChem, converted to PDBQT format using PyMOL software (Kim et al., 2016; Yuan et al., 2017). Blind docking was performed using AutoDock Vina protein-size-adjusted grid boxes, and top-ranked poses were selected based on binding affinity for visualization and interaction analysis (Trott and Olson, 2010; Yuan et al., 2017). Protein-ligand interactions were analyzed using PLIP, and key non-covalent interactions (hydrogen bonds, hydrophobic interactions, π-π interactions, and salt bridges) were visualized in PyMOL (Salentin et al., 2015; Yuan et al., 2017).

### 2.11. Molecular dynamics simulation of RutA-EE2 and RutA-warfarin sodium complexes

The top-ranked RutA-EE2 and RutA-warfarin sodium docking complexes were subjected to MD simulation in GROMACS within the Galaxy Europe environment using the AMBER99SB force field (Gu et al., 2023b; Hashem et al., 2025; Heine et al., 2018; Kim et al., 2010; Valton et al., 2006). Each protein-ligand system was solvated in a triclinic TIP3P water box, neutralized with counterions, equilibrated, and simulated for 100 ps to assess complex stability. Trajectories were analysed for Root Mean Square Deviation (RMSD), Root Mean Square Fluctuations (RMSF), and radius of gyration (Rg), and energy components (Potential energy). These parameters were used to evaluate structural stability and convergence of the RutA-ligand complex during simulation.

### 2.12. Statistical analysis

Experiments were performed in triplicates unless stated otherwise, and data were expressed as mean ± standard deviation. Growth values were analysed as log CFU mL^-1^.

Statistical analysis was performed using IBM SPSS Statistics v29 (George and Mallery, 2024). Time-course growth data were analysed using repeated-measures general linear model (GLM). For substrate growth and PAβN assays, Time and Treatment were used as within-subject factors. For induction assays, Time and Induction treatment were used as within-subject factors. Estimated marginal means were compared using Bonferroni-adjusted pairwise comparisons.

Warfarin sodium depletion was analysed by fitting residual concentration to a first-order kinetic model. The apparent rate constant was calculated from the slope of the ln (C) versus time plot, and the half-life was calculated as t_1/2_ = 0.693 k^-1^. Model fit was assessed using the regression coefficient.

Differences were considered statistically significant at p < 0.05.

## 3. Results

### 3.1. Isolation and primary screening of SS02 on EE2 and warfarin sodium

Strain SS02 was recovered from hospital sludge after four successive enrichment cycles in MSM supplemented with EE2 (10 mg L^-1^) as the sole carbon source (Fig. 1A). After enrichment, visible bacterial colonies appeared on EE2-MSM agar (Fig. 1B). SS02 also showed visible growth on MSM agar containing warfarin sodium (20 mg L^-1^) as the sole added carbon source (Fig. 1B). In addition, SS02 produced visible growth on Luria agar (LA), confirming viability and culturability on a heterotrophic medium (Fig. 1B) Growth in liquid MSM supplemented with warfarin sodium (20 mg L^-1^) further confirmed that SS02 remained viable and active under warfarin sodium exposure, supporting its compatibility with a structurally distinct pharmaceutical substrate. No visible growth was observed in substrate-free MSM control. Abiotic and heat-killed controls were used to exclude non-biological changes during subsequent assays. Morphological and biochemical characterization identified strain SS02 as a rod-shaped, Gram-negative bacterium (Table S4), with mean length of 2.243 µm and mean breadth of 0.921 µm obtained from scanning electron micrograph (SEM) of bacterial strain SS02 (Fig. 1C)

**Fig. 1.**
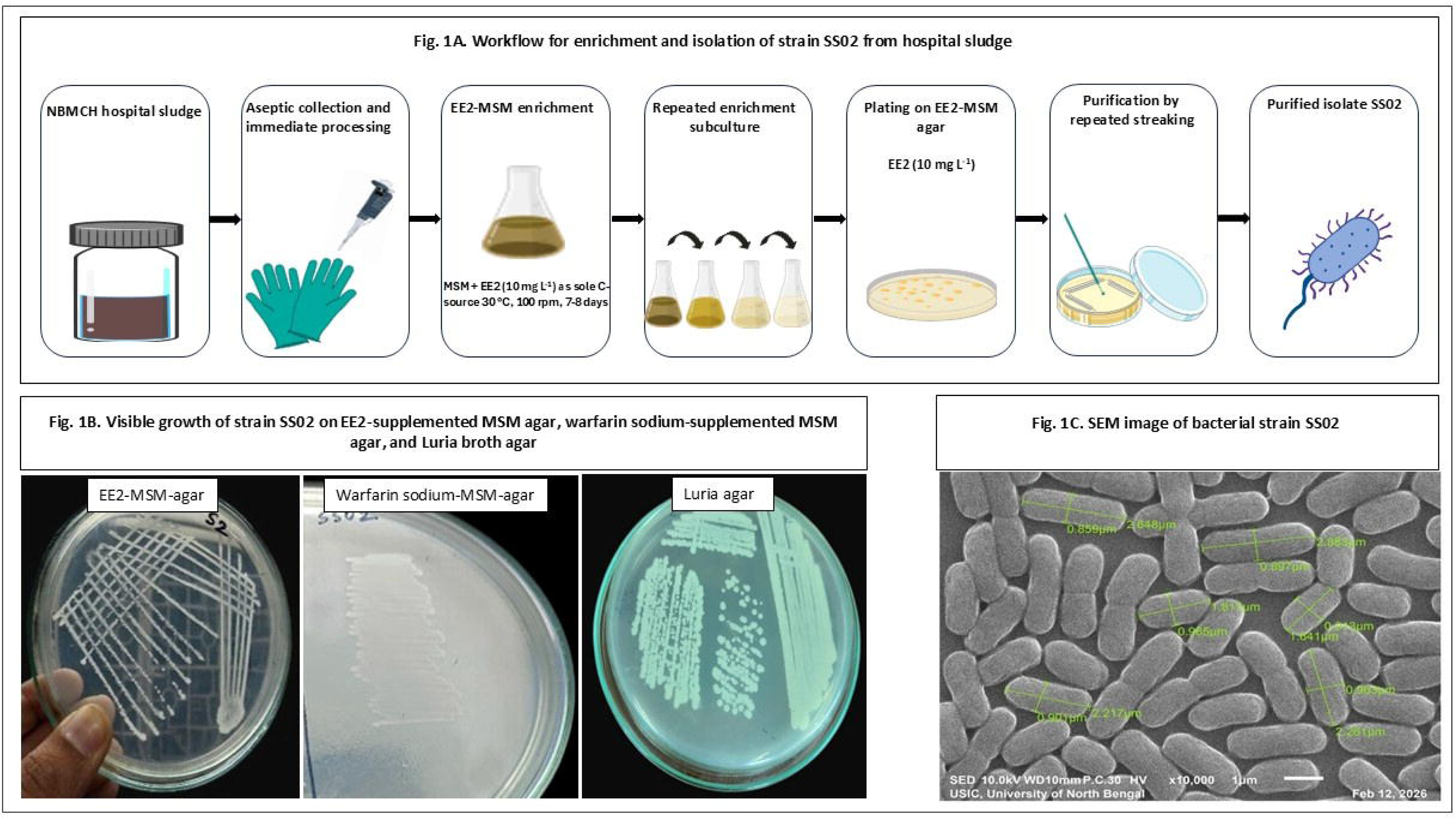
Isolation and primary screening of strain SS02 on EE2 and warfarin sodium. **(A)** Workflow showing enrichment of hospital sludge in EE2-supplemented MSM, serial enrichment, and isolation of strain SS02. **(B)** Visible growth of SS02 on MSM agar supplemented with EE2 as the sole carbon source, warfarin sodium as sole carbon source, and Luria agar (heterotrophic medium) **(C)** SEM image of bacterial strain SS02.

### 3.2. Growth of SS02 on EE2, warfarin sodium, and combined EE2 + warfarin sodium

Strain SS02 grew in all three MSM conditions and showed a clear time-dependent rise in viable counts. Initial counts were about 3.2 to 3.5 log CFU mL^-1^ at 0 h. By 12 h, viable cell counts increased to about 5.3 to 5.6 log CFU mL^-1^. By 24 h, growth reached about 6.4 log CFU mL^-1^ in warfarin sodium, about 6.6 log CFU mL^-1^ in EE2 and about 6.7 to 6.8 log CFU mL^-1^ in the combined condition.

Repeated-measures analysis confirmed a significant effect of time on SS02 growth, showing progressive increase in viable counts across all substrate conditions. The combined-substrate condition showed numerically higher growth during the early to mid-phase, especially between 12 and 48 h. However, Bonferroni-adjusted pairwise comparisons did not detect significant differences among EE2, warfarin sodium, and combined EE2 + warfarin sodium treatments. By 48 to 60 h, all treatments approached similar final viable counts near 7.0 log CFU mL^-1^. These results indicate that SS02 is compatible with EE2, warfarin sodium, and mixed EE2 + warfarin sodium exposure, showing a trend toward faster early growth rather than a statistically distinct endpoint response (Fig.2A).

**Fig. 2.**
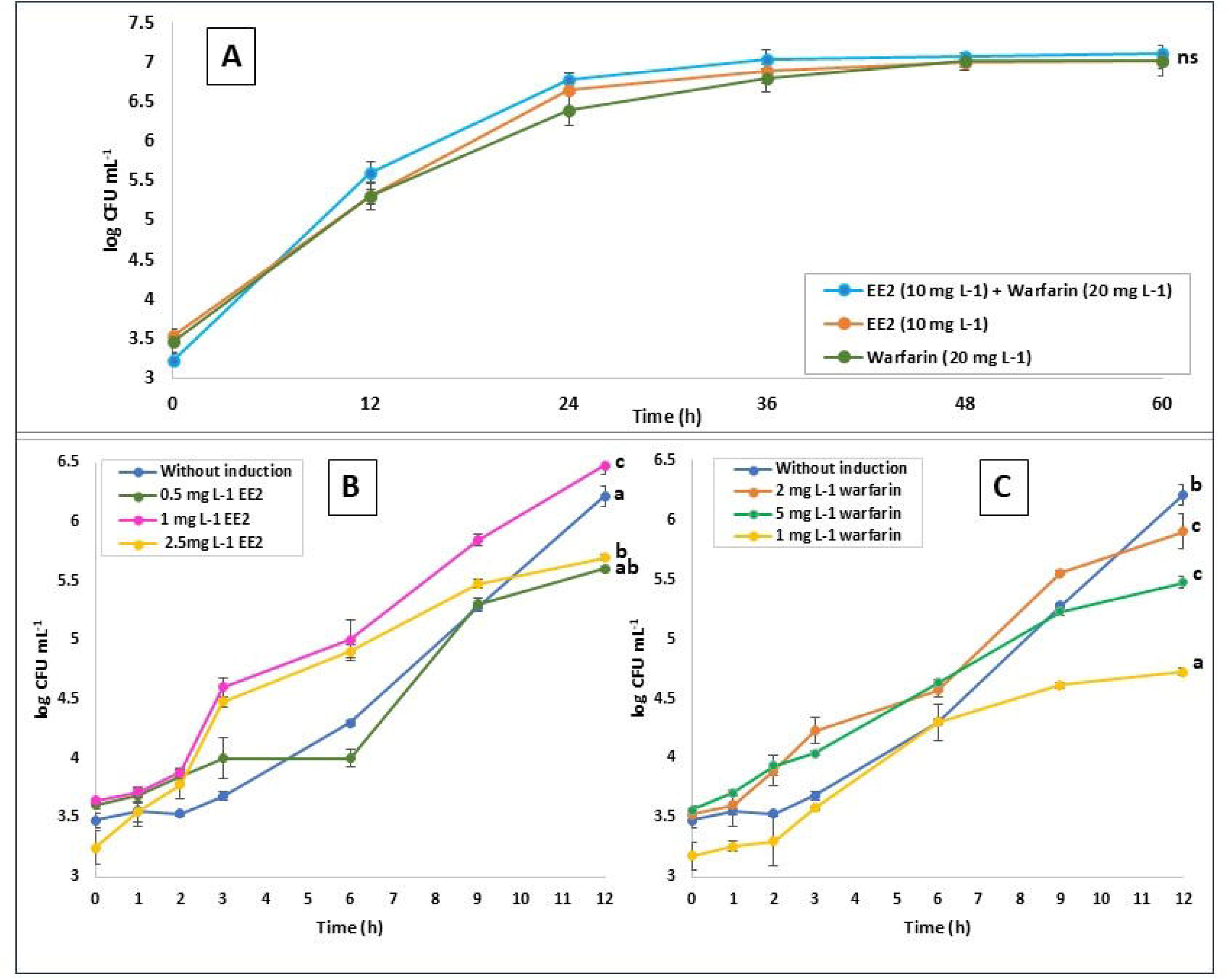
Growth and induction responses of *Klebsiella pneumoniae* SS02 under EE2 and warfarin sodium exposure. **(A)** Growth of SS02 in MSM supplemented with EE2, warfarin sodium, or combined EE2 + warfarin sodium. **(B)** Growth response after pre-exposure to different concentrations of EE2, followed by transfer to EE2-supplemented MSM. **(C)** Growth response after pre-exposure to different concentrations of warfarin sodium, followed by transfer to EE2-supplemented MSM. Viable counts were monitored as log CFU mL□¹ over time. Values represent mean ± SD of triplicate experiments. Repeated-measures analysis showed a significant time-dependent increase in viable counts. In panel A, Bonferroni-adjusted pairwise comparisons did not detect significant differences among substrate treatments. In panel B and C, induction treatment significantly altered the growth response. Different letters indicate significant differences among treatments based on Bonferroni-adjusted pairwise comparisons, p < 0.05. Same letters indicate no significant difference. ns indicates no significant pairwise difference among substrate treatments.

### 3.3. EE2 and warfarin sodium induce early growth response in SS02

Pre-exposure to EE2 and warfarin sodium influenced the subsequent growth of SS02 in EE2-supplemented MSM. In the EE2 induction assay, repeated-measures analysis showed significant effects of time, induction treatment, and Time x Induction interaction, indicating altered growth kinetics of SS02. The non-induced control increased gradually from 3.47 log CFU mL^-1^ at 0 h to 6.21 log CFU mL^-1^ at 12 h. Among the EE2 induction treatments, 1 mg L^-1^ EE2 produced the strongest response and differed significantly from the non-induced control, p= 0.019, and from 2.5 mg L^-1^ EE2, p = 0.024. The 0.5 mg L^-1^ treatment did not differ significantly from the control, indicating concentration dependent induction with strongest response at 1 mg L^-1^ (Fig. 2B).

Warfarin sodium pre-exposure also enhanced subsequent EE2-supported growth, indicating cross-induction. Repeated-measures analysis showed significant effects of time, induction treatment, and Time x Induction interaction. The strongest response was observed after 2 mg L^-1^ warfarin sodium induction, followed by 5 mg L^-1^. Both treatments produced significantly higher responses than the non-induced control, p = 0.013 and p = 0.011, respectively. The 1 mg L^-1^ warfarin sodium treatment showed the weakest response and was significantly lower than the control, 2 mg L^-1^ and 5 mg L^-1^ treatments. At 12 h, viable counts reached about 6.07, 5.44, and 4.69 log CFU mL^-1^ after induction with 2, 5, and 1 mg L^-1^ warfarin sodium, 5.44 log CFU mL^-1^ after 5 mg L^-1^ induction, and 4.69 log CFU mL^-1^ after 1 mg L^-1^ warfarin sodium, respectively (Fig.2C).

Overall, both EE2 and warfarin sodium pre-exposure improved early EE2-supported growth of SS02. EE2 induction was strongest at 1 mg L^-1^, while warfarin sodium induction was strongest at 2 mg L^-1^. These results support overlapping substrate-responsive machinery rather than simple passive tolerance.

### 3.4. UV-Vis tracking shows delayed warfarin sodium utilization during combined EE2 + warfarin sodium exposure

Warfarin sodium showed a linear UV-Vis response at 309 nm across 0.0025 to 0.025 mg mL^-1^. The calibration equation was y= 55.291x – 0.0477 with R^2^ = 0.9988, confirming high linearity for quantification of residual warfarin sodium in culture supernatants (Fig. S1).

The calibration curve was used to quantify residual warfarin sodium in the medium. In warfarin sodium-only MSM, viable counts increased from about 3.4 log CFU mL^-1^ at 0 h to about 6.8 log CFU mL^-1^ at 36 h. During the same period, residual warfarin sodium decreased from about 18.3 mg L^-1^ to about 13.0 mg L^-1^. This indicates growth-associated warfarin sodium depletion (Fig. 3A).

**Fig. 3.**
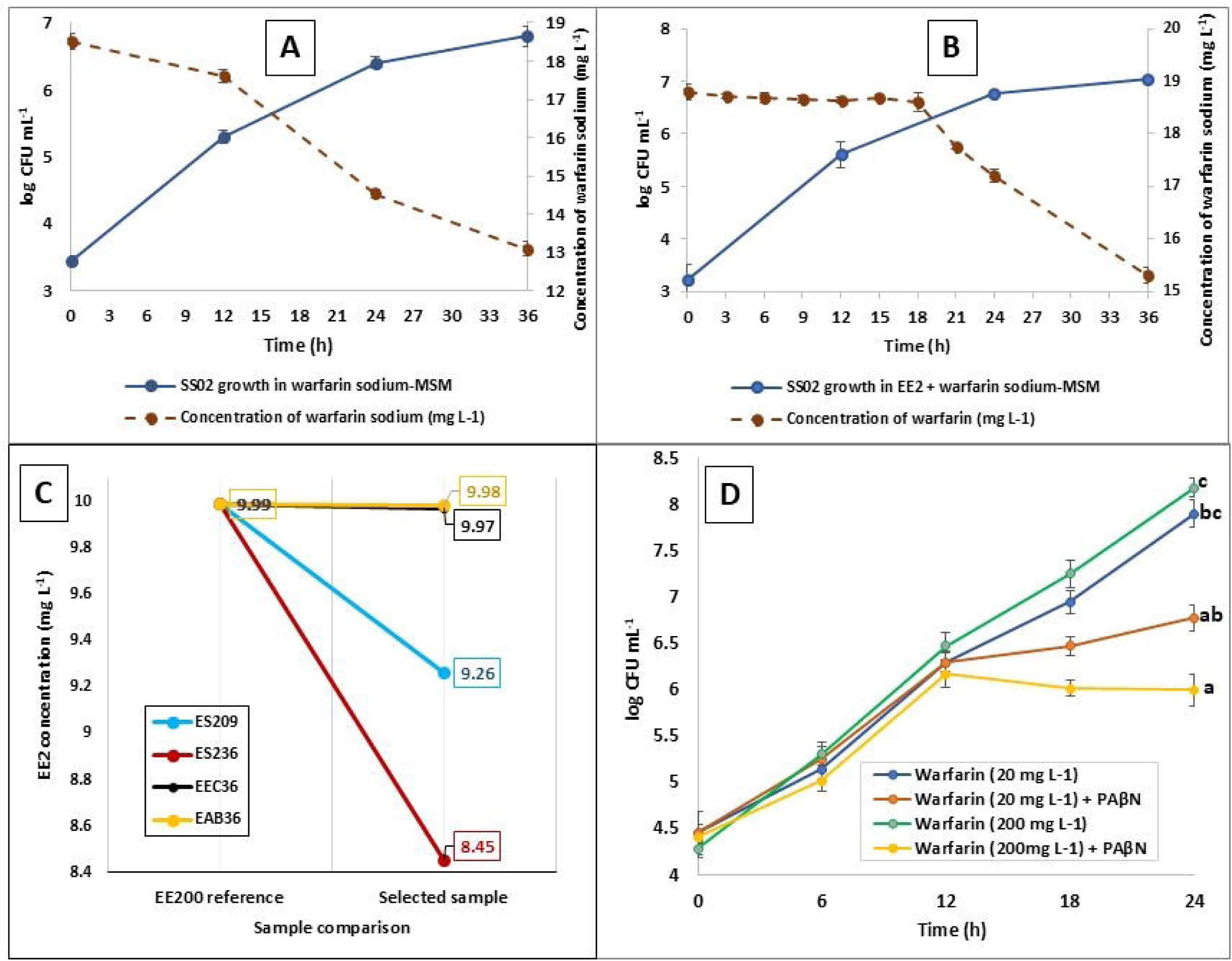
Warfarin sodium utilization and efflux-associated tolerance in *Klebsiella pneumoniae* SS02. **(A)** Growth of SS02 and residual warfarin sodium concentration in MSM containing warfarin sodium (20 mg L□¹) as the sole carbon source **(B)** Growth of SS02 and residual warfarin sodium concentration in combined EE2 (10 mg L^-1^) + warfarin sodium (20 mg L□¹) MSM. **(C)** Reference-to-sample change in parent EE2 concentration based on targeted UPLC-MS/MS/MRM analysis. EE200 represents the EE2 0 h reference sample; ES209 and ES236 represents SS02 culture supernatants at 9 and 36 h, respectively; EEC36 represents *E. coli* K12 control at 36 h; and EAB36 represents the abiotic control at 36 h. **(D)** Effect of PAβN on growth of SS02 under warfarin sodium stress. Viable counts are expressed as log CFU mL□¹, and residual warfarin sodium was estimated by UV-Vis absorbance at 309 nm. Values represent mean ± SD of triplicate experiments. Repeated-measures analysis showed significant effects of time, treatment, and Time x Treatment interaction in panel D. Different letters indicate significant differences among treatments based on Bonferroni-adjusted pairwise comparisons, p < 0.05. Same letters indicate no significant difference.

Using these residual concentration values, first order kinetics was observed for warfarin sodium removal. The ln(C) versus time plot showed a linear fit with the equation y= −0.0102x + 2.9431 and R^2^ = 0.9565, giving an apparent first-order rate constant k = 0.0102 h^-1^ and half-life t_1/2_ = 0.693 k^-1^ = 67.94 h (Fig. S2).

In combined EE2 + warfarin sodium medium, SS02 also showed strong growth, increasing from about 3.2 log CFU mL^-1^ at 0 h to about 7.04 log CFU mL^-1^ at 36 h. However, warfarin sodium concentration remained nearly stable up to 18 h, and decreased sharply only after 21 h, reaching about 15.3 mg L^-1^ by 36 h (Fig. 3B). This supports delayed warfarin sodium utilization under combined exposure, where early growth is likely supported by EE2-linked response, followed by later warfarin sodium transformation.

### 3.4. Targeted UPLC-MS/MS analysis of EE2

Targeted UHPLC-MS/MS/MRM analysis detected the parent EE2 signal at approximately 0.91 min. The EE2 pure-compound calibration showed good linearity, with R^2^ = 0.996 (Fig.S3). The LOD and LOQ were 2.78 and 8.41 mg L^-1^, respectively. EE2 0 h reference sample contained 0.99 µg mL^-1^ EE2. Compared with 0 h sample, EE2 in 9 h and 36 h supernatants decreased to 9.25 and 8.45 mg L^-1^ respectively, corresponding to approximately 7.4% and 15.45% depletion respectively. In contrast, abiotic and *E. coli* K12 controls remained close to the reference level. The reference-to-sample change plot supports condition-dependent depletion of EE2, with the strongest reduction observed at 36 h sample (Fig.3C).

### 3.5 PA***β***N reduces growth under warfarin sodium stress, supporting efflux-assisted tolerance

SS02 tolerated both 20 mg L^-1^ and 200 mg L^-1^ warfarin sodium in absence of efflux inhibitor. At 20 mg L^-1^ warfarin sodium, viable counts increased from about 4.45 to 7.9 log CFU mL^-1^ by 24 h. Under elevated warfarin sodium stress, 200 mg L^-1^, SS02 also sustained growth, reaching about 8.2 log CFU mL^-1^ at 24 h (Fig.3D).

PAβN reduced growth under both warfarin sodium conditions. At 20 mg L^-1^ warfarin sodium, PAβN-treated cultures reached only about 6.8 log CFU mL^-1^ at 24 h, compared with about 7.9 log CFU mL^-1^ in the untreated culture. The effect was stronger at 200 mg L^-1^, where PAβN-treated cultures remained near 6.0 log CFU mL^-1^ at 18 h and 24 h, while untreated cultures continued to increase (Fig.3D). This indicates that efflux activity contributes to warfarin sodium tolerance in SS02.

Repeated-measures analysis confirmed significant effects of time, treatment, and Time x Treatment interaction on SS02 growth. Using Greenhouse-Geisser correction, the time effect was significant, F = 1574.804, p < 0.001, partial η^2^ = 0.999. The treatment effect was also significant, F= 113.331, p = 0.001, partial η^2^ = 0.983. The Time x Treatment interaction was significant, F= 29.013, p = 0.005, partial η^2^ = 0.936, indicating that PAβN altered the growth trajectory over time. Bonferroni-adjusted comparisons showed that the 200 mg L^-1^ warfarin sodium condition without PAβN differed significantly from both PAβN-treated conditions, supporting an efflux-associated contribution to growth under warfarin sodium stress.

### 3.6. Whole genome sequencing, phylogenomic placement and biosafety screening of SS02

Whole-genome sequencing of strain SS02 generated a draft genome of 5.54 Mbp. The genome was annotated using the NCBI Prokaryotic Genome Annotation Pipeline and RAST, which provided the basis for downstream functional screening of genes associated with EE2 response, aromatic metabolism, regulation, and efflux-associated tolerance.

Whole genome based taxonomic placement identified strain SS02 as *Klebsiella pneumoniae*. Phylogenomic analysis using the Type Strain Genome Server placed SS02 within the *Klebsiella pneumoniae* lineage (Fig. S4). This placement was independently reproduced using the KBase phylogenomics workflow (Fig.S5), and both analyses showed concordant topology. The agreement between TYGS and KBase supported the taxonomic assignment of SS02 as *Klebsiella pneumoniae*.

The whole genome sequence was submitted to the National Center for Biotechnology Information (NCBI) database (GenBank No.: JAWLIH000000000) (Benson, 2004).

In addition to taxonomic placement, genome-based screening was used to assess the biosafety profile of SS02 (Note S1).

### 3.7. Genome mining identifies RutA, aromatic-processing genes, and efflux determinants

Genome mining of strain SS02 identified a set of genes potentially associated with EE2 and warfarin sodium response. Functional annotation using PGAP, RAST and eggNOG mapper revealed candidate enzymes involved in oxidative transformation, aromatic compound processing, redox metabolism, and efflux-associated tolerance (Table S5).

RutA was identified as a key candidate protein in the genome. Its annotation as a flavin-dependent monooxygenase-like protein supports its possible role in early oxidative activation of pharmaceutical substrates. Because both EE2 and warfarin sodium contain ring-based structures requiring oxidative transformation before downstream processing. RutA was selected as the proposed oxidative entry enzyme in the pathway model.

The genome also contained several genes linked to aromatic ring-processing metabolism. These include HpaB/HpaC-like hydroxylase components, oxidoreductases, dehydrogenases, hydrolases, and genes associated with *cat*, *pca*, *ben*, and β-ketoadipate-linked pathways. These modules support the possibility that oxidized EE2- and warfarin sodium derived intermediates are further processed through aromatic degradation and carbon-funnelling routes (Table S5).

Efflux-related genes were also detected, including RND-type multidrug efflux determinants such as *mdtABC* and *acrAB-tolC*. Their presence supports the efflux-tolerance components of the model and agrees with the PAβN assay, where efflux inhibition reduced growth under warfarin sodium stress. Therefore, genome mining supports a combined metabolic-defence strategy in SS02, involving RutA-linked oxidative entry, downstream aromatic processing, and efflux-assisted tolerance (Table S5).

### 3.8. Conserved domain analysis supports RutA as a flavin-dependent oxidative entry protein

Conserved domain analysis of RutA from strain SS02 showed that the protein contains domains consistent with flavin-dependent monooxygenase and oxidoreductase activity. The strongest hit was identified as RutA, a pyrimidine utilization protein A, spanning almost the full length, from amino acid 1 to 355, with an E-value of 0. This supports the annotation of the protein as a RutA-type monooxygenase.

Additional domain hits supported the same functional assignment. The protein showed similarity to alkanesulfonate monooxygenase, luciferase-like monooxygenase, nitrilotriacetate monooxygenase, and flavin-dependent oxidoreductase families. These domains are commonly associated with flavin-mediated oxidation reactions. The presence of the SsuD-like flavin-utilizing monooxygenase domain further supports the involvement of RutA in oxidative catalysis.

The conserved domain profile therefore supports RutA as a putative flavin-dependent oxidative entry protein in SS02 (Table 1).

**Table 1.**
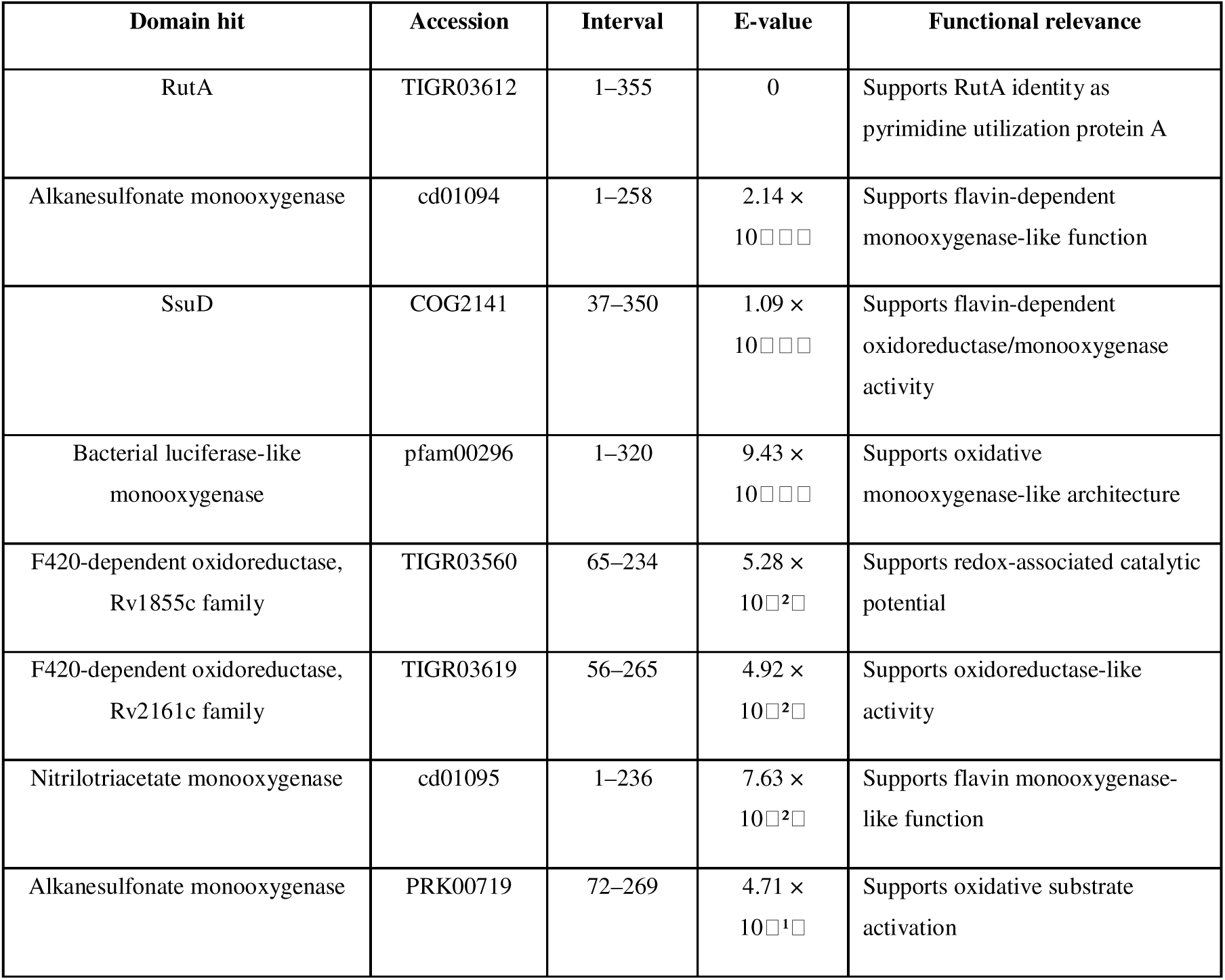
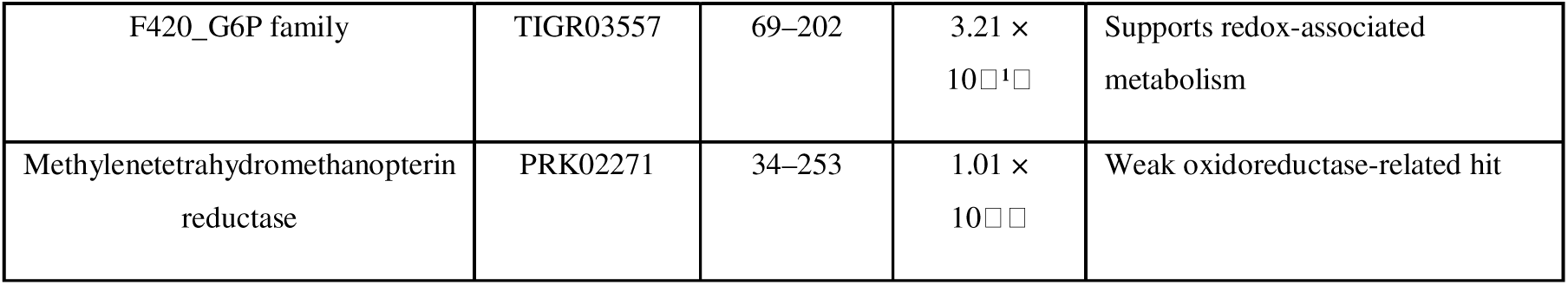
Conserved domain features of RutA from *Klebsiella pneumoniae* SS02.

### 3.9. STRING network resolves linked entry, redox-processing, aromatic-catabolic, and efflux modules

STRING-based protein-protein interaction analysis resolved the candidate SS02 proteins into five functionally coherent modules (Fig.4). K-means clustering separated the network into modules linked to Rut-associated entry reactions, Hpa-associated redox processing, β-ketoadipate-linked aromatic metabolism, downstream carbon funnelling, and efflux-associated tolerance. This topology supported a coordinated response system rather than isolated gene-level functions.

**Fig. 4.**
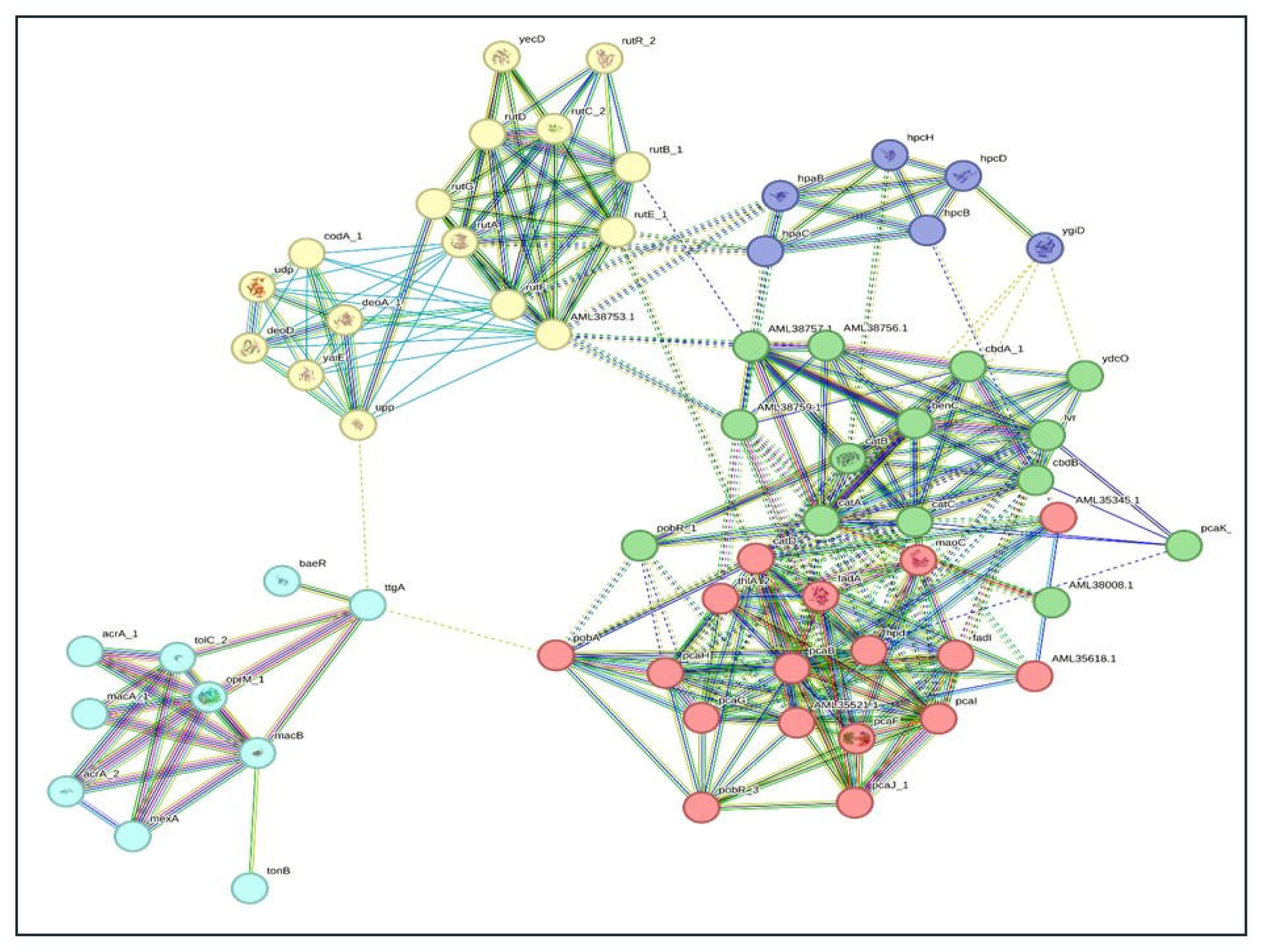
STRING PPI network of warfarin sodium and EE2 response module in strain SS02, clustered into functional groups. The network highlights an entry oxidation cluster (*rut* genes), an aromatic hydroxylation module (*hpa* genes), a dense aromatic catabolic core (*ben*, *cat* and *pca* genes of the β-ketoadipate route), and an efflux and envelope defense module (AcrAB-TolC, MexAB-OprM, MacAB, with BaeR). Nodes represent proteins and edges represents high-confidence functional associations inferred from STRING evidence channels.

The Rut module formed the primary entry node of the network. RutA was centrally positioned and closely associated with proteins encoded by *rutB*, *rutC*, *rutF*, and *rutG*. This arrangement supports the proposed role of RutA as a shared oxidative gateway for EE2 and warfarin sodium response. The Hpa module, comprising proteins encoded by *hpaB*, *hpaC*, and *hpaD*, connected the entry module with downstream metabolism, suggesting redox-supported transformation of early intermediates.

A densely connected β-ketoadipate-associated core formed the major downstream metabolic hub. Proteins encoded by *pca*, *cat*, and *ben* genes were grouped in this core, indicating a strong aromatic-catabolic module. This supports the proposed conversion of EE2- and warfarin sodium-derived aromatic intermediates through ring-cleavage and central carbon-funneling routes. Downstream proteins, including FadA and associated enzymes, extended this processing toward acetyl-CoA and succinyl-CoA pools, suggesting links with energy generation and redox balance.

A distinct efflux-associated module contained AcrAB-TolC, MexAB-OprM, and MacAB, with BaeR positioned as a regulatory node. This module was separated from the metabolic core but retained limited bridging interactions, suggesting stress-responsive, linkage between intracellular transformation and efflux-mediated defense. This agrees with the PAβN inhibition assay, where efflux inhibition reduced SS02 growth under elevated warfarin sodium stress.

Overall, the STRING network supported a two-arm operational model in SS02. In the first arm, EE2 and warfarin sodium are proposed to enter through RutA-linked oxidative processing and are progressively funnelled through Hpa and β-ketoadipate-associated aromatic metabolism. In the second arm, efflux systems support survival by exporting toxic substrates or intermediates and maintaining intracellular stability.

### 3.10. Hypothesized EE2-warfarin sodium response pathway places RutA at the shared entry point

Based on the combined phenotypic, genomic, network, and structural evidence, a hypothesized EE2-warfarin sodium response pathway was constructed for strain SS02 (Fig.5). The model placed RutA at the shared entry point of the proposed pathway because RutA was detected in both EE2- and warfarin sodium-associated gene inventories and showed conserved domain features consistent with flavin-dependent oxidative activity.

**Fig. 5.**
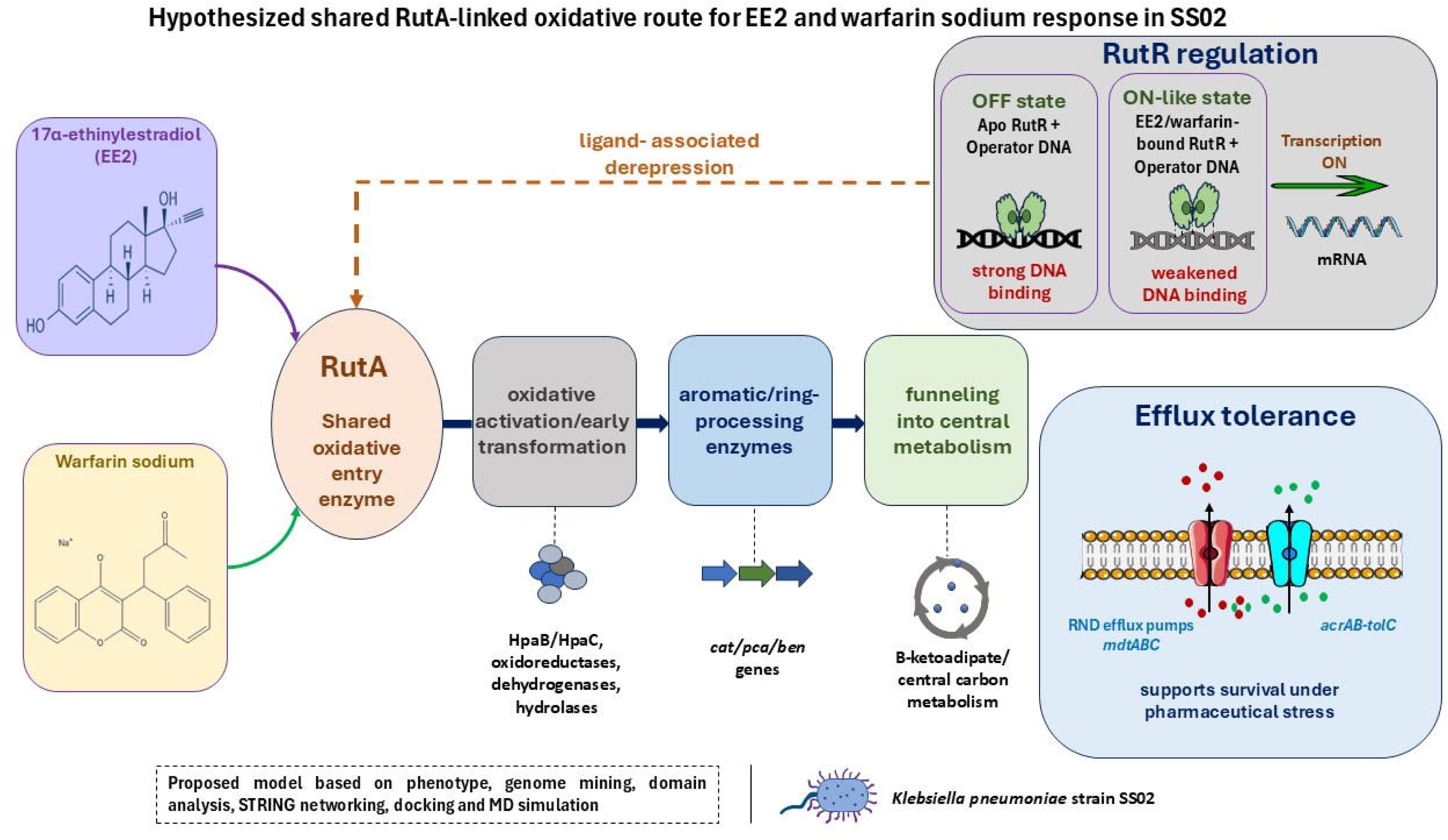
Proposed RutA-linked oxidative response model for EE2 and warfarin sodium adaptation in *Klebsiella pneumoniae* SS02. EE2 and warfarin sodium are proposed to enter through RutA-mediated oxidative activation, followed by redox processing, aromatic/ring transformation, and β-ketoadipate-linked carbon funneling. RutR regulation is proposed to shift from strong DNA binding in the apo state to weakened DNA binding in ligand-bound ON-like states. RND-type efflux systems, including mdtABC and acrAB-tolC, support pharmaceutical stress tolerance. The model integrates phenotype, genome mining, STRING networking, docking, and MD simulation.

In the proposed model, EE2 and warfarin sodium converge at RutA for early oxidative activation. This step is followed by redox-supported transformation through HpaB/HpaC, oxidoreductases, dehydrogenases, and hydrolases. The resulting aromatic intermediates are proposed to enter ring-processing routes involving *cat, pca, ben*, and β-ketoadipate-associated genes. These downstream modules provide a plausible route for funnelling transformed substrate into central carbon metabolism.

The model also included a RutR-linked regulatory arm. Apo RutR bound to operator DNA was interpreted as the OFF-state model, where strong DNA binding maintains transcriptional repression. In contrast, EE2- or warfarin sodium-bound RutR showed weakened DNA engagement and was interpreted as an ON-like state. The reduced RutR-DNA interaction was proposed to permit transcriptional activation of the RutR-linked response, represented by mRNA production in the model (Fig.5). This regulatory component provides a structural explanation for the induction and cross-induction response observed after EE2 or warfarin sodium pre-exposure.

Efflux determinants, including mdtABC and acrAB-tolC, were placed as tolerance modules rather than catabolic enzymes. This placement was supported by the PAβN inhibition assay, where efflux inhibition reduced growth under warfarin sodium stress. Thus, SS02 appears to combine oxidative metabolism with efflux-assisted tolerance during pharmaceutical exposure.

Overall, the pathway supports a two-component response model. The first component involves RutA-linked oxidative entry followed by aromatic processing and carbon funneling. The second component involves RutR-associated regulation and efflux-mediated defense. This model explains how SS02 grows on EE2 and warfarin sodium and why both substrates appear to activate overlapping response machinery (Fig.5).

### 3.11 RutR docking supports ligand-associated derepression of the Rut response

RutR-operator DNA docking showed a clear reduction in protein-DNA interface contact after ligand binding. The apo RutR-DNA complex, representing the OFF state, showed the strongest DNA engagement, with 100 protein atoms within 4 Å of DNA, and 102 DNA atoms were within 4 Å of the protein. In contrast, the EE2-bound RutR-DNA model showed reduced interface contact, with 71 protein atoms and 76 DNA atoms were within 4 Å (Fig. 6A,6B). When residue-level contacts were selected using byres, the apo model showed 227 protein atoms and 479 DNA atoms at the interface, whereas the EE2-bound RutR-DNA showed 138 protein atoms and 351 DNA atoms, and warfarin sodium-bound RutR-DNA model showed 153 protein atoms and 351 DNA atoms. This reduction indicates weaker RutR-operator engagement after EE2 binding (Table S6).

**Fig. 6.**
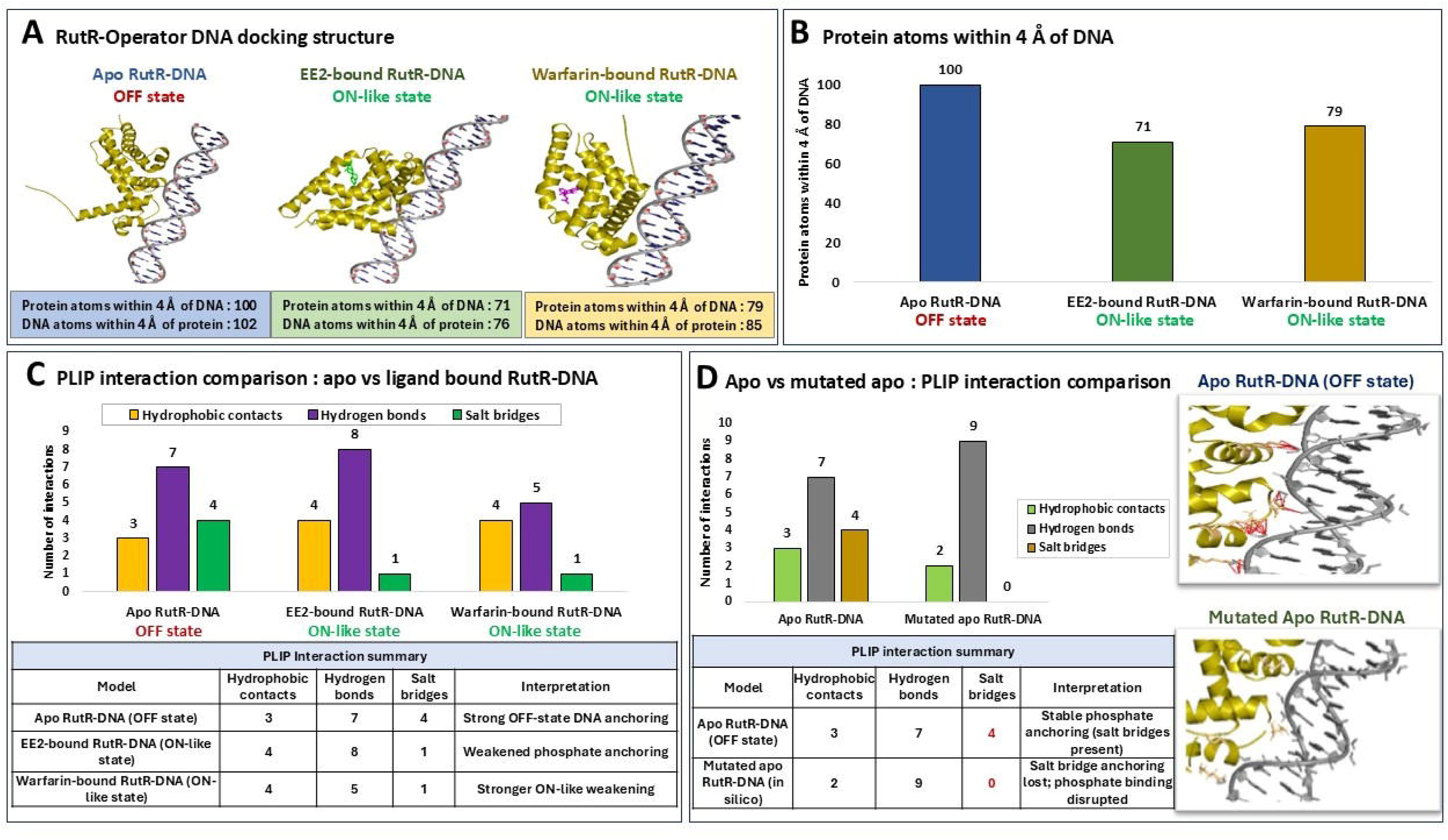
RutR-operator DNA docking and PLIP-based interaction analysis. **(A)** Docked structures of apo RutR-DNA, EE2-bound RutR-DNA, and warfarin sodium-bound RutR-DNA, showing stronger DNA engagement in the apo OFF state and reduced DNA contact in ligand-bound ON-like states. **(B)** Protein atoms within 4 Å of DNA decreased from 100 in apo RutR-DNA to 71 and 79 in EE2-bound and warfarin sodium-bound RutR-DNA models, respectively. **(C)** PLIP comparison showed reduced salt-bridge anchoring in ligand-bound RutR-DNA models. **(D)** In silico mutation of selected DNA-contacting residues abolished apo RutR-DNA salt bridges, supporting the role of phosphate-backbone anchoring in the OFF-state model.

To validate the structural basis of this OFF-to-ON-like transition, the protein-DNA interfaces of apo and ligand bound RutR models were further analysed using PLIP. This analysis showed that the apo RutR-DNA retained the strongest DNA-binding profile, with 3 hydrophobic contacts, 7 hydrogen bonds, and 4 salt bridges. In contrast, the EE2- and warfarin sodium-bound models retained only 1 salt bridge each, indicating loss of major phosphate-backbone anchoring in the ligand-bound states (Fig. 6C).

To further assess the contribution of DNA-contacting residues, selected PLIP-identified RutR interface residues were mutated in silico and reanalysed (Table S7). The apo model showed 4 salt bridges involving His37, Arg47, Lys52, and Lys113. After mutation, all salt bridges were lost, although hydrogen bonds were redistributed. The mutated apo RutR model retained 2 hydrophobic contacts and showed 9 hydrogen bonds, but no salt bridges were detected (Fig.6D). This indicates that mutation of key interface residues did not abolish all local contacts, but specifically disrupted the strong electrostatic anchoring between RutR and the DNA phosphate backbone. The increased hydrogen-bond count in the mutant likely reflects contact redistribution after side-chain alteration, not stronger DNA repression. Therefore, the mutation result supports the PLIP-based conclusion that salt bridge-mediates phosphate anchoring is a key feature of the apo RutR OFF-state complex.

Together, interface-count analysis, PLIP profiling, and in silico mutation support the same trend: apo RutR forms a strong DNA-bound OFF-state complex, while ligand-bound RutR shows weakened DNA engagement mainly through loss of salt bridge-mediated phosphate anchoring (Fig. 6D). This supports ligand-associated derepression model for the Rut response.

### 3.12. Docking supports RutA interaction with both EE2 and warfarin sodium

Genome mining suggested that EE2 and warfarin sodium may require an initial oxidative activation step before downstream aromatic processing. Molecular docking was therefore performed with candidate oxidative and efflux-associated proteins to identify plausible substrate-binding targets. Among the tested proteins, RutA emerged as the strongest common candidate for both substrates.

EE2 showed strong binding to RutA, with a binding affinity of −9.9 kcal mol^-1^. The top-ranked pose was positioned within an internal RutA pocket. Warfarin sodium also showed strong binding to RutA, with a binding affinity of −9.1 kcal mol^-1^, indicating a comparable interaction with the same candidate oxidative scaffold (Fig.S6).

PLIP analysis supported stable placement of both substrates in the RutA pocket. EE2 formed multiple hydrophobic contacts with Phe6, Leu46, Met48, Val117, Ala187, Phe205, Phe207, and Leu328, along with two hydrogen bonds involving Trp120 and Leu240. Warfarin sodium also formed extensive hydrophobic contacts with Phe6, Leu46, Val117, Ala187, Phe205, Leu240, Asn286, and Leu328, together with three hydrogen bonds involving Asn115, Trp120, and Asn286. The shared involvement of residues such as Phe6, Leu46, Val117, Ala187, Phe205, and Leu328 suggest that EE2 and warfarin sodium interact with overlapping RutA pocket regions (Fig.S7).

Docking of warfarin sodium with efflux-associated proteins also produced plausible binding affinities, including MdtB (−7.7 kcal mol^-1^), MdtC (−7.6 kcal mol^-1^), and AcrAB (−7.8 kcal mol^-1^) (Fig.S6). These results support the efflux-tolerance module and agree with the PAβN inhibition assay, where efflux inhibition reduced growth under warfarin sodium stress.

### 3.13. MD simulation supports stability of RutA-ligand complexes

To evaluate binding stability beyond docking, the top ranked RutA-EE2 and RutA-warfarin sodium complexes were subjected to molecular dynamics simulation in GROMACS. The overall structural stability, residue flexibility, compactness and energetic stability were assessed using RMSD, RMSF, radius of gyration and potential energy.

RMSD analysis showed rapid stabilization of both complexes after the initial equilibration phase. The RutA-EE2 complex maintained a backbone RMSD around 0.10 to 0.12 nm for most of the trajectory, with only a mild drift of 0.13 to 0.14 nm near the end. The RutA-Warfarin sodium complex likewise remained stable, with backbone RMSD around 0.09 to 0.13 nm across the 100 ps trajectory and only a small shift around 40-55 ps, before a stable plateau was reached (Fig.S8).

RMSF profiles remained low for most residues in both the complexes, typically around 0.04 to 0.08 nm in RutA-EE2 and around 0.04 to 0.07 nm in RutA-Warfarin sodium complex, with only small peaks in localized loop regions (Fig.S8).

Radius of gyration values remained stable near 2.0 nm throughout both simulations, indicating retained compactness without unfolding or major expansion (Fig.S8).

RutA-EE2 complex showed that the potential energy stabilized quickly initially and remained stable around approximately −5.1 x 10^5^ kJ mol^-1^ with small fluctuations, indicating the system is well-equilibrated. The potential energy in case of RutA-warfarin sodium complex showed rapid convergence during the start and remained stable throughout the run, with a small fluctuation around 5.21 x 10^5^ kJ mol^-1^, indicating a well-equilibrated system (Fig.S8).

## 4. Discussion

Hospital-sludge represents a chemically complex environment containing mixtures of pharmaceuticals, redox-active compounds, and antimicrobial stressors. Such environments can select for metabolically flexible and stress-tolerant microorganisms (Patel et al., 2019; Santos et al., 2010). The research has provided proof for the fact that *Klebsiella pneumoniae* SS02 isolated from the hospital sludge shows a co-ordinated response towards structurally diverse pharmaceuticals such as 17α-ethinylestradiol (EE2) and warfarin sodium. Instead of indicating completely separate metabolic pathways, the physiological, chemical, and genomic data has proved that these pharmaceuticals utilize similar adaptive machinery, leading to coupled metabolic-defense mechanism.

An important finding in this study was the different performance of substrates when subjected to a mixed treatment. Although both EE2 and warfarin sodium were able to sustain the growth when used separately, the slower depletion rate of warfarin sodium in the presence of EE2 along with the observed depletion of EE2 itself implies a sequence or a hierarchy in substrate usage. This indicates that the oxidation pathways related to EE2 could be stimulated first, followed by the increased oxidation of warfarin sodium. This agrees with reports that microbial estrogen transformation commonly involves oxidative reactions followed by assimilation or co-metabolic routing into central metabolism (Anes et al., 2015; Chen et al., 2018; Hashem et al., 2025; Ibero et al., 2020).

Analytically, targeted UHPLC-MS/MS revealed the dependence of reduction of parent EE2 on conditions, whereas warfarin sodium exhibited growth-linked reduction following a first-order kinetic trend. The apparent-first-order depletion pattern is consistent with biodegradation under non-saturating substrate conditions (Buchan et al., 2000). While transformation products could not be detected, the observed parallel trends of growth and substrate reduction suggest the activity of microorganisms. The lack of an additive effect of growth, when subjected to both compounds strengthen the possibility of common metabolic pathways.

Genome-based studies have provided a mechanistic rationale behind such an interpretation. The presence of RutA as a flavin monooxygenase enzyme family member and the presence of Hpa-related reductases along with β-ketoadipate-related aromatic degradation genes suggest that there is a possible oxidative pathway entry and funnelling model for EE2 and warfarin sodium metabolism. RutA is classically linked with pyrimidine metabolism, but its flavin-dependent oxygenation chemistry provides a plausible biochemical basis for early oxidative handling of ring-containing substrates (Gu et al., 2023a; Hashem et al., 2025; Heine et al., 2018; Valton et al., 2006). Nevertheless, the function of RutA still remains speculative and should be taken only as a candidate for the oxidative pathway entry point and not as an enzyme.

The regulatory aspect of this response can be explained through RutR model simulations where the destabilization of DNA binding due to ligand interaction serves as a possible way of gene derepression. EE2-bound and warfarin sodium-bound RutR-DNA models showed reduced protein-DNA interface contact (Nguyen Le Minh et al., 2015). This serves as a potential structural basis for the induction and cross induction, but needs to be confirmed experimentally (EMSA or transcriptomics).

At the same time, efflux system mediated tolerance is another important component. Lowering of growth as a result of exposure to PAβN especially under increased warfarin sodium exposure is a direct indication of the role of the RND-type efflux systems in regulating intracellular homeostasis. Since RND systems mediate Gram-negative tolerance to diverse toxicants (Delmar et al., 2014; Sun et al., 2014), and MdtABC is linked with bulky compound export and BaeSR-regulated envelope stress (Baranova and Nikaido, 2002; Kim et al., 2020), efflux likely acts as an accessory tolerance mechanism rather than a primary degradation route. Identification of the acrAB-tolC and mdtABC systems through genome mining serves as additional evidence supporting this point. All these observations fit perfectly into the framework of the idea that metabolic modification alone is not enough when dealing with increased contamination.

Overall, the findings suggest that a two-armed model of adaptation is applicable to SS02. One arm includes oxidation and aromatic metabolism which can be initiated through the activity of RutA-dependent reactions in association with β-ketoadipate metabolism. Such coupling is relevant in contaminated habitats, where hydrophobic or partially oxidized intermediates may impose membrane and redox stress (Ramos et al., 2002; Segura et al., 2005; Shi et al., 2019). The β-ketoadipate pathway provides a central route for aerobic aromatic assimilation and links catechol- and protocatechuate-derived intermediates to central metabolism (Buchan et al., 2000; Harwood and Parales, 1996; Romero-Silva et al., 2013). The other arm includes tolerance by means of efflux and helps the bacteria survive under the pressure of pharmaceuticals.

With regards to the environment, the combination of these mechanisms for adaptation could prove very useful in sludge-related microbial populations in which pharmaceutical compounds occur as complex mixtures at various concentrations. This could have an effect on the survival and degradation of these contaminants in the process of wastewater treatment. It must be mentioned, however, that the concentration of the chemicals used in this experiment is more akin to enrichment level rather than environmental conditions.

Several limitations should be kept in mind. No transformation products of EE2 and warfarin sodium were reported, and thus the proposed pathway cannot be confirmed metabolomically. The model for oxidative entry via RutA lacks experimental confirmation as it has been suggested solely on the basis of functional annotation, domain prediction, network analysis, and modeling. This model explains grows on both substrates, the absence of a strongly additive endpoint response under combined exposure, warfarin sodium-linked cross induction of EE2-supported growth, and delayed warfarin sodium depletion during mixed exposure (Buchan et al., 2000; Harwood and Parales, 1996). Analogously, the regulation mechanism that involves RutR requires confirmation of its existence by means of protein expression profiling or DNA binding experiments.

However, despite these weaknesses, this research represents an integration of mechanistic insights into bacterial adaptation to combined drug exposure. The integration of substrate metabolism, regulatory control, and efflux mechanism contributes to increasing our knowledge about how bacteria survive and operate in drug-contaminated environments. This information is crucial for predicting the behaviour of contaminants and developing efficient biological approaches to wastewater purification.

## 5. Conclusion

This study shows that a *Klebsiella pneumoniae* strain (SS02) isolated from hospital sludge is able to adapt to the mixed exposure of structurally different pharmaceuticals, 17α-ethinylestradiol (EE2) and warfarin sodium. It was demonstrated through growth kinetics, induction profile, and analytical data that these compounds use similar adaptive response strategies and do not undergo different metabolic pathways. The quantitative degradation of the parent EE2 and growth-dependent decrease of warfarin sodium along with retarded consumption by mixed exposure were observed.

Based on genome-based and structural studies, the mechanisms involve an oxidative entrance mechanism, which is probably linked to RutA and associated with the processing of aromatics. Alongside this, efflux plays an important role in the resistance to pharmaceuticals. This information provides strong evidence for a two-branch model for adaptation that involves both metabolism and efflux.

Despite the fact that the framework is based on converging evidence at the level of phenotype, analysis, genetics, and in silico predictions, it is still a hypothesis-driven framework lacking metabolite-resolved data and enzymatic verification. Therefore, future research should focus on obtaining metabolite-resolved profiling, testing of the candidate enzymes, and regulatory validations in order to validate the model.

On the whole, this paper presents a mechanistic view of the possible way of survival of the environmental bacteria under exposure to a mixture of pharmaceuticals. Such metabolic-defensive strategies are expected to impact contaminant fate in sludge affected ecosystems and can be used for biological-based wastewater treatments.

## AUTHOR CONTRIBUTION

**Shilpa Sinha:** Investigation, Methodology, Data curation, Formal analysis, Validation, Writing – original draft, Writing – review and editing.

**Partha Barman:** Investigation, Methodology, Formal analysis, Data curation, Visualization, Writing – review and editing.

**Debopriya Haldar:** Investigation, Data curation, Formal analysis.

**Ranadhir Chakraborty:** Conceptualization, Methodology, Supervision, Formal analysis, Data curation, Project administration, Resources, Visualization, Writing – review and editing.

## ACKOWLEDGEMENTS

The authors sincerely acknowledge Miss Riddhi Chakraborty, IIT Madras, India, for her assistance in method calibration during the initial phase of warfarin sodium quantification using the UV–Vis method.

## DATA AVAILABILITY STATEMENT

the sequencing data pertinent to this study is available at the NCBI database under the BioProject ID: PRJNA1026355, BioSample ID: SAMN37738782, carrying GenBank accession ID: JAWLIH000000000, while sequencer-generated raw reads are available from the NCBI Sequence Reads Archives, bearing the SRA accession ID: SRR27015460.

## DECLARATION OF GENERATIVE AI USE

The authors declare the use of ChatGPT v5.4 (OpenAI) during manuscript preparation, to assist in drafting narrative text. No AI tool was used to generate, manipulate, or analyse experimental data or results. All AI-generated content was critically reviewed to ensure accuracy and scientific integrity.

---

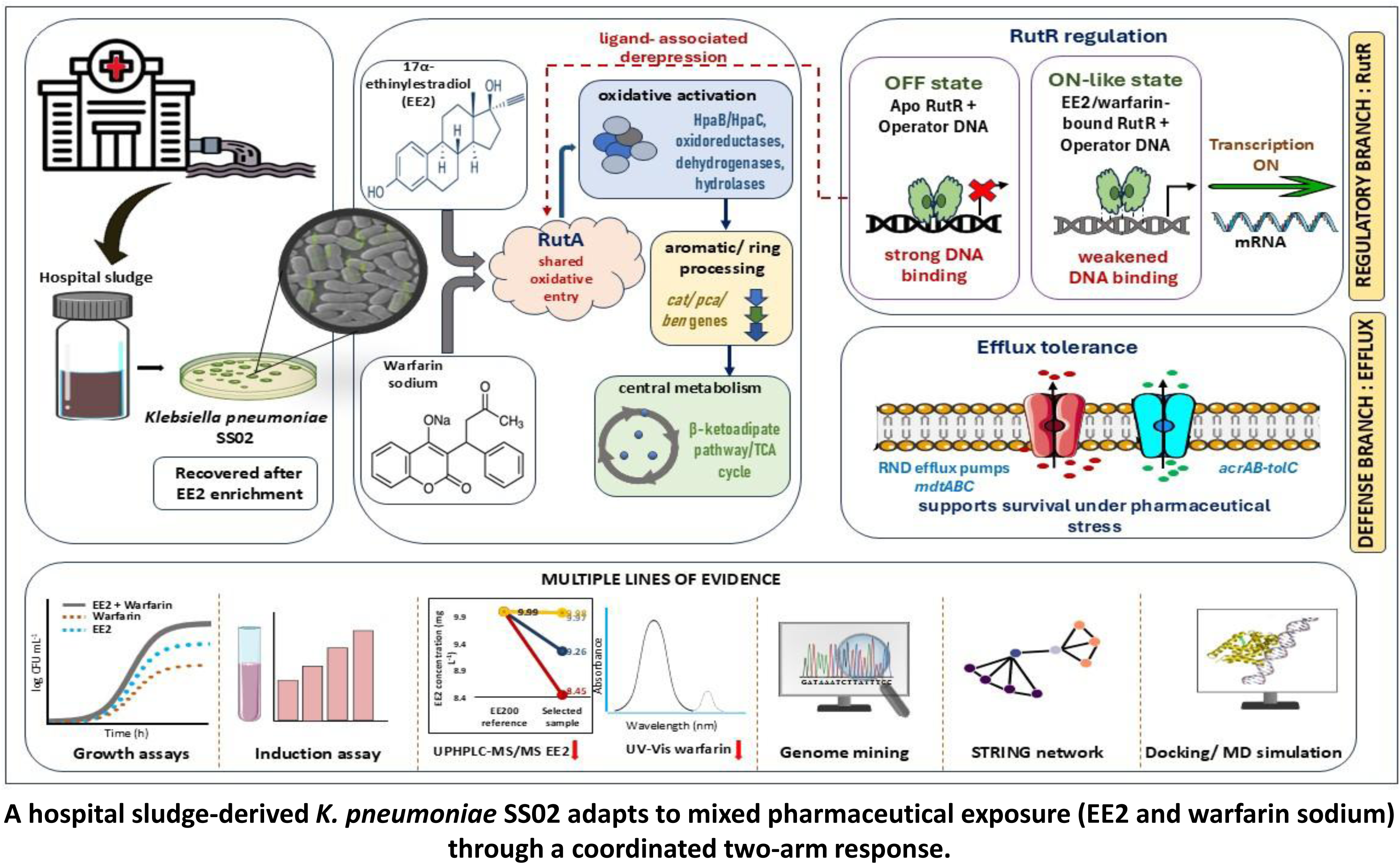

## References

Alexander, Martin., 1999. Biodegradation and bioremediation. Academic Press.

Aneja, K.R., 2018. Experiments in microbiology, plant pathology, tissue culture and microbial biotechnology, 5th ed.

Anes, J., McCusker, M.P., Fanning, S., Martins, M., 2015. The ins and outs of RND efflux pumps in Escherichia coli. Front. Microbiol. 6. 10.3389/fmicb.2015.00587

Arkin, A.P., Stevens, R.L., Cottingham, R.W., Maslov, S., Henry, C.S., Dehal, P., Ware, D., Perez, F., Harris, N.L., Canon, S., Sneddon, M.W., Henderson, M.L., Riehl, W.J., Gunter, D., Murphy-Olson, D., Chan, S., Kamimura, R.T., Brettin, T.S., Meyer, F., Chivian, D., Weston, D.J., Glass, E.M., Davison, B.H., Kumari, S., Allen, B.H., Baumohl, J., Best, A.A., Bowen, B., Brenner, S.E., Bun, C.C., Chandonia, J.-M., Chia, J.-M., Colasanti, R., Conrad, N., Davis, J.J., DeJongh, M., Devoid, S., Dietrich, E., Drake, M.M., Dubchak, I., Edirisinghe, J.N., Fang, G., Faria, J.P., Frybarger, P.M., Gerlach, W., Gerstein, M., Gurtowski, J., Haun, H.L., He, F., Jain, R., Joachimiak, M.P., Keegan, K.P., Kondo, S., Kumar, V., Land, M.L., Mills, M., Novichkov, P., Oh, T., Olsen, G.J., Olson, B., Parrello, B., Pasternak, S., Pearson, E., Poon, S.S., Price, G.A., Ramakrishnan, S., Ranjan, P., Ronald, P.C., Schatz, M.C., Seaver, S.M.D., Shukla, M., Sutormin, R.A., Syed, M.H., Thomason, J., Tintle, N.L., Wang, D., Xia, F., Yoo, H., Yoo, S., 2016. The DOE Systems Biology Knowledgebase (KBase). 10.1101/096354

Aziz, R.K., Bartels, D., Best, A.A., DeJongh, M., Disz, T., Edwards, R.A., Formsma, K., Gerdes, S., Glass, E.M., Kubal, M., Meyer, F., Olsen, G.J., Olson, R., Osterman, A.L., Overbeek, R.A., McNeil, L.K., Paarmann, D., Paczian, T., Parrello, B., Pusch, G.D., Reich, C., Stevens, R., Vassieva, O., Vonstein, V., Wilke, A., Zagnitko, O., 2008. The RAST Server: Rapid Annotations using Subsystems Technology. BMC Genomics 9, 75. 10.1186/1471-2164-9-75

Babhair, S.A., Tariq, M., Al-Badr, A.A., 1985. Warfarin. pp. 423–452. 10.1016/S0099-5428(08)60587-0

Baranova, N., Nikaido, H., 2002. The BaeSR Two-Component Regulatory System Activates Transcription of the *yegMNOB* (*mdtABCD*) Transporter Gene Cluster in *Escherichia coli* and Increases Its Resistance to Novobiocin and Deoxycholate. J. Bacteriol. 184, 4168–4176. 10.1128/JB.184.15.4168-4176.2002

Bergey, D. Hendricks., 1994. Bergey’s manual of determinative bacteriology.

Buchan, A., Collier, L.S., Neidle, E.L., Moran, M.A., 2000. Key Aromatic-Ring-Cleaving Enzyme, Protocatechuate 3,4-Dioxygenase, in the Ecologically Important Marine Roseobacter Lineage. Appl. Environ. Microbiol. 66, 4662–4672. 10.1128/AEM.66.11.4662-4672.2000

Cantalapiedra, C.P., Hernández-Plaza, A., Letunic, I., Bork, P., Huerta-Cepas, J., 2021. eggNOG-mapper v2: Functional Annotation, Orthology Assignments, and Domain Prediction at the Metagenomic Scale. Mol. Biol. Evol. 38, 5825–5829. 10.1093/molbev/msab293

Chatterjee, S., Barman, P., Barman, C., Majumdar, S., Chakraborty, R., 2024. Multimodal cadmium resistance and its regulatory networking in Pseudomonas aeruginosa strain CD3. Sci. Rep. 14, 31689. 10.1038/s41598-024-80754-y

Chen, Y.-L., Fu, H.-Y., Lee, T.-H., Shih, C.-J., Huang, L., Wang, Y.-S., Ismail, W., Chiang, Y.-R., 2018. Estrogen Degraders and Estrogen Degradation Pathway Identified in an Activated Sludge. Appl. Environ. Microbiol. 84. 10.1128/AEM.00001-18

Chen, Y.-L., Yu, C.-P., Lee, T.-H., Goh, K.-S., Chu, K.-H., Wang, P.-H., Ismail, W., Shih, C.-J., Chiang, Y.-R., 2017. Biochemical Mechanisms and Catabolic Enzymes Involved in Bacterial Estrogen Degradation Pathways. Cell Chem. Biol. 24, 712–724.e7. 10.1016/j.chembiol.2017.05.012

Delmar, J.A., Su, C.-C., Yu, E.W., 2014. Bacterial Multidrug Efflux Transporters. Annu. Rev. Biophys. 43, 93–117. 10.1146/annurev-biophys-051013-022855

Fernández, I., Santos, A., Cancela, M.L., Laizé, V., Gavaia, P.J., 2014. Warfarin, a potential pollutant in aquatic environment acting through Pxr signaling pathway and γ-glutamyl carboxylation of vitamin K-dependent proteins. Environmental Pollution 194, 86–95. 10.1016/j.envpol.2014.07.015

George, D., Mallery, P., 2024. IBM SPSS Statistics 29 Step by Step A Simple Guide and Reference. Routledge, New York. 10.4324/9781032622156

Gómez-Canela, C., Barata, C., Lacorte, S., 2014. Occurrence, elimination, and risk of anticoagulant rodenticides and drugs during wastewater treatment. Environmental Science and Pollution Research 21, 7194–7203. 10.1007/s11356-014-2714-1

Gu, Y., Li, T., Yin, C.-F., Zhou, N.-Y., 2023a. Elucidation of the coumarin degradation by Pseudomonas sp. strain NyZ480. J. Hazard. Mater. 457, 131802. 10.1016/j.jhazmat.2023.131802

Gu, Y., Li, T., Zhou, N.-Y., 2023b. Redundant and scattered genetic determinants for coumarin biodegradation in *Pseudomonas* sp. strain NyZ480. Appl. Environ. Microbiol. 89. 10.1128/aem.01109-23

Harwood, C.S., Parales, R.E., 1996. The β-Ketoadipate pathway and the biology of self-identity. Annu. Rev. Microbiol. 50, 553–590. 10.1146/annurev.micro.50.1.553

Hashem, J.S., Ismail, W., Chiang, Y.-R., Bekhit, A.A., 2025. Diversity of estrogen biodegradation pathways and application in environmental bioremediation. Front. Microbiol. 16. 10.3389/fmicb.2025.1630636

Heine, T., Van Berkel, W.J.H., Gassner, G., Van Pée, K.-H., Tischler, D., 2018. Two-Component FAD-Dependent Monooxygenases: Current Knowledge and Biotechnological Opportunities. Biology (Basel). 7, 42. 10.3390/biology7030042

Hsiao, T., Chen, Y., Meng, M., Chuang, M., Horinouchi, M., Hayashi, T., Wang, P., Chiang, Y., 2021. Mechanistic and phylogenetic insights into actinobacteria□mediated oestrogen biodegradation in urban estuarine sediments. Microb. Biotechnol. 14, 1212–1227. 10.1111/1751-7915.13798

Ibero, J., Galán, B., Rivero-Buceta, V., García, J.L., 2020. Unraveling the 17β-Estradiol Degradation Pathway in Novosphingobium tardaugens NBRC 16725. Front. Microbiol. 11. 10.3389/fmicb.2020.588300

Joss, A., Zabczynski, S., Göbel, A., Hoffmann, B., Löffler, D., McArdell, C.S., Ternes, T.A., Thomsen, A., Siegrist, H., 2006. Biological degradation of pharmaceuticals in municipal wastewater treatment: Proposing a classification scheme. Water Res. 40, 1686–1696. 10.1016/j.watres.2006.02.014

Kim, H., Kim, S., Kim, D., Yoon, S.H., 2020. A single amino acid substitution in aromatic hydroxylase (HpaB) of Escherichia coli alters substrate specificity of the structural isomers of hydroxyphenylacetate. BMC Microbiol. 20, 109. 10.1186/s12866-020-01798-4

Kim, K.-S., Pelton, J.G., Inwood, W.B., Andersen, U., Kustu, S., Wemmer, D.E., 2010. The Rut Pathway for Pyrimidine Degradation: Novel Chemistry and Toxicity Problems. J. Bacteriol. 192, 4089–4102. 10.1128/JB.00201-10

Kim, S., Thiessen, P.A., Bolton, E.E., Chen, J., Fu, G., Gindulyte, A., Han, L., He, J., He, S., Shoemaker, B.A., Wang, J., Yu, B., Zhang, J., Bryant, S.H., 2016. PubChem Substance and Compound databases. Nucleic Acids Res. 44, D1202–D1213. 10.1093/nar/gkv951

Meier-Kolthoff, J.P., Göker, M., 2019. TYGS is an automated high-throughput platform for state-of-the-art genome-based taxonomy. Nat. Commun. 10, 2182. 10.1038/s41467-019-10210-3

Morris, G.M., Huey, R., Olson, A.J., 2008. Using AutoDock for Ligand□Receptor Docking. Curr. Protoc. Bioinformatics 24. 10.1002/0471250953.bi0814s24

Nguyen Le Minh, P., de Cima, S., Bervoets, I., Maes, D., Rubio, V., Charlier, D., 2015. Ligand binding specificity of RutR, a member of the TetR family of transcription regulators in *Escherichia coli*. FEBS Open Bio 5, 76–84. 10.1016/j.fob.2015.01.002

Patel, M., Kumar, R., Kishor, K., Mlsna, T., Pittman, C.U., Mohan, D., 2019. Pharmaceuticals of Emerging Concern in Aquatic Systems: Chemistry, Occurrence, Effects, and Removal Methods. Chem. Rev. 119, 3510–3673. 10.1021/acs.chemrev.8b00299

Ramos, J.L., Duque, E., Gallegos, M.-T., Godoy, P., Ramos-González, M.I., Rojas, A., Terán, W., Segura, A., 2002. Mechanisms of Solvent Tolerance in Gram-Negative Bacteria. Annu. Rev. Microbiol. 56, 743–768. 10.1146/annurev.micro.56.012302.161038

Romero-Silva, M.J., Méndez, V., Agulló, L., Seeger, M., 2013. Genomic and Functional Analyses of the Gentisate and Protocatechuate Ring-Cleavage Pathways and Related 3-Hydroxybenzoate and 4-Hydroxybenzoate Peripheral Pathways in Burkholderia xenovorans LB400. PLoS One 8, e56038. 10.1371/journal.pone.0056038

Salentin, S., Schreiber, S., Haupt, V.J., Adasme, M.F., Schroeder, M., 2015. PLIP: fully automated protein–ligand interaction profiler. Nucleic Acids Res. 43, W443–W447. 10.1093/nar/gkv315

Santos, L.H.M.L.M., Araújo, A.N., Fachini, A., Pena, A., Delerue-Matos, C., Montenegro, M.C.B.S.M., 2010. Ecotoxicological aspects related to the presence of pharmaceuticals in the aquatic environment. J. Hazard. Mater. 175, 45–95. 10.1016/j.jhazmat.2009.10.100

Segura, A., Godoy, P., van Dillewijn, P., Hurtado, A., Arroyo, N., Santacruz, S., Ramos, J.-L., 2005. Proteomic Analysis Reveals the Participation of Energy- and Stress-Related Proteins in the Response of *Pseudomonas putida* DOT-T1E to Toluene. J. Bacteriol. 187, 5937–5945. 10.1128/JB.187.17.5937-5945.2005

Shi, X., Chen, M., Yu, Z., Bell, J.M., Wang, H., Forrester, I., Villarreal, H., Jakana, J., Du, D., Luisi, B.F., Ludtke, S.J., Wang, Z., 2019. In situ structure and assembly of the multidrug efflux pump AcrAB-TolC. Nat. Commun. 10, 2635. 10.1038/s41467-019-10512-6

Sun, J., Deng, Z., Yan, A., 2014. Bacterial multidrug efflux pumps: Mechanisms, physiology and pharmacological exploitations. Biochem. Biophys. Res. Commun. 453, 254–267. 10.1016/j.bbrc.2014.05.090

Szklarczyk, D., Gable, A.L., Lyon, D., Junge, A., Wyder, S., Huerta-Cepas, J., Simonovic, M., Doncheva, N.T., Morris, J.H., Bork, P., Jensen, L.J., von Mering, C., 2019. STRING v11: protein–protein association networks with increased coverage, supporting functional discovery in genome-wide experimental datasets. Nucleic Acids Res. 47, D607–D613. 10.1093/nar/gky1131

Tang, Z., Liu, Z., Wang, H., Dang, Z., Liu, Y., 2021a. A review of 17α-ethynylestradiol (EE2) in surface water across 32 countries: Sources, concentrations, and potential estrogenic effects. J. Environ. Manage. 292, 112804. 10.1016/j.jenvman.2021.112804

Tang, Z., Liu, Z., Wang, H., Dang, Z., Liu, Y., 2021b. Occurrence and removal of 17α-ethynylestradiol (EE2) in municipal wastewater treatment plants: Current status and challenges. Chemosphere 271, 129551. 10.1016/j.chemosphere.2021.129551

Tatusova, T., DiCuccio, M., Badretdin, A., Chetvernin, V., Nawrocki, E.P., Zaslavsky, L., Lomsadze, A., Pruitt, K.D., Borodovsky, M., Ostell, J., 2016. NCBI prokaryotic genome annotation pipeline. Nucleic Acids Res. 44, 6614–6624. 10.1093/nar/gkw569

Tian, K., Meng, Q., Li, S., Chang, M., Meng, F., Yu, Y., Li, H., Qiu, Q., Shao, J., Huo, H., 2022. Mechanism of 17β-estradiol degradation by Rhodococcus equi via the 4,5-seco pathway and its key genes. Environmental Pollution 312, 120021. 10.1016/j.envpol.2022.120021

Trott, O., Olson, A.J., 2010. AutoDock Vina: Improving the speed and accuracy of docking with a new scoring function, efficient optimization, and multithreading. J. Comput. Chem. 31, 455–461. 10.1002/jcc.21334

Valton, J., Fontecave, M., Douki, T., Kendrew, S.G., Nivière, V., 2006. An Aromatic Hydroxylation Reaction Catalyzed by a Two-component FMN-dependent Monooxygenase. Journal of Biological Chemistry 281, 27–35. 10.1074/jbc.M506146200

van Dijk, M., 2006. Information-driven protein-DNA docking using HADDOCK: it is a matter of flexibility. Nucleic Acids Res. 34, 3317–3325. 10.1093/nar/gkl412

Varadi, M., Anyango, S., Deshpande, M., Nair, S., Natassia, C., Yordanova, G., Yuan, D., Stroe, O., Wood, G., Laydon, A., Žídek, A., Green, T., Tunyasuvunakool, K., Petersen, S., Jumper, J., Clancy, E., Green, R., Vora, A., Lutfi, M., Figurnov, M., Cowie, A., Hobbs, N., Kohli, P., Kleywegt, G., Birney, E., Hassabis, D., Velankar, S., 2022. AlphaFold Protein Structure Database: massively expanding the structural coverage of protein-sequence space with high-accuracy models. Nucleic Acids Res. 50, D439–D444. 10.1093/nar/gkab1061

Wang, J., Chitsaz, F., Derbyshire, M.K., Gonzales, N.R., Gwadz, M., Lu, S., Marchler, G.H., Song, J.S., Thanki, N., Yamashita, R.A., Yang, M., Zhang, D., Zheng, C., Lanczycki, C.J., Marchler-Bauer, A., 2023. The conserved domain database in 2023. Nucleic Acids Res. 51, D384–D388. 10.1093/nar/gkac1096

Yuan, S., Chan, H.C.S., Hu, Z., 2017. Using <SCP>PyMOL<SCP> as a platform for computational drug design. WIREs Computational Molecular Science 7. 10.1002/wcms.1298

