## Supplemental Table for "A Two-Arm Metabolic-Efflux Adaptation Framework in *Klebsiella pneumoniae* under Mixed Pharmaceutical Exposure: *rutA*-Linked Oxidative Entry and *rutR*-Associated Regulation"

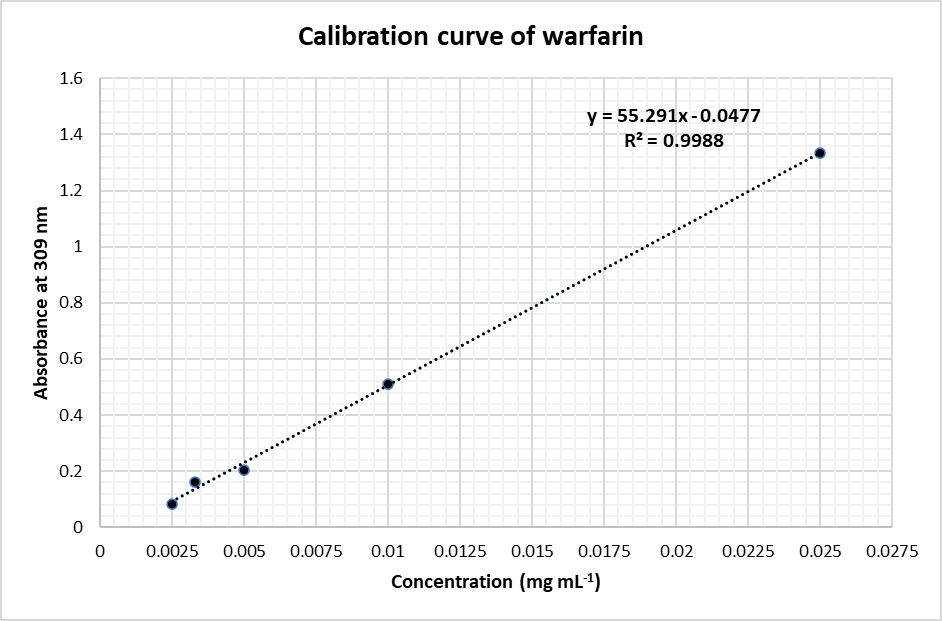

**Fig. S1.** UV-Vis calibration curve of warfarin sodium in MSM at 309 nm (0.0025 – 0.025 mg mL^-1^), showing linearity (y = 55.291x – 0.0477; R^2^ = 0.9988).

Apparent first order degradation kinetics of warfarin sodium by strain SS02

ln C

| Time (h) | SS02 [Warfarin (mg/L) (ln C)] |
| --- | --- |
| 0 | 2.9178 |
| 12 | 2.8687 |
| 24 | 2.6764 |
| 36 | 2.5726 |

**Fig. S2.** Apparent first order depletion kinetics of warfarin sodium by SS02, fitted from residual concentration data. The fitted regression showed k = 0.0102 h^-1^ and R^2^ = 0.956.

Calibration curve for EE2 quantification by UHPLC-MS/MS/MRM

| Concentration of EE2 (mg L-1) | Mean Area |
| --- | --- |
| 3.12 | 316.76 |
| 6.25 | 685.98 |
| 12.5 | 1725.63 |
| 25 | 3128.35 |
| 50 | 6383.13 |

**Fig. S3.** Calibration curve for EE2 quantification by UHPLC-MS/MS/MRM. The calibration curve was generated using EE2 pure compound standards, with mean peak area plotted against concentration. The regression equation was **y = 128.73x – 45.975, with R^2^ = 0.9985**

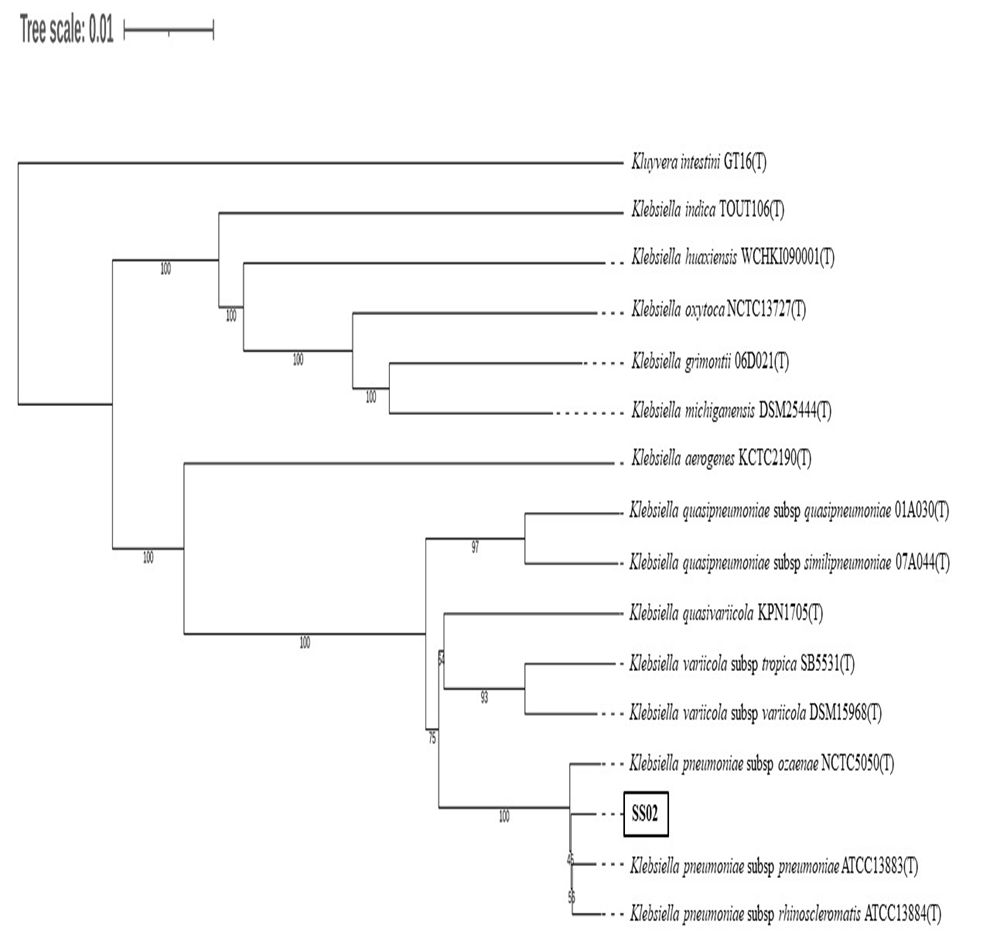

**Fig. S4.** Whole genome-based phylogenetic tree using TYGS server, showing the taxonomic placement of strain SS02 within the *Klebsiella pneumoniae* clad. Bootstrap supports values are shown at the internal nodes, and the scale bar indicates substitution per site.

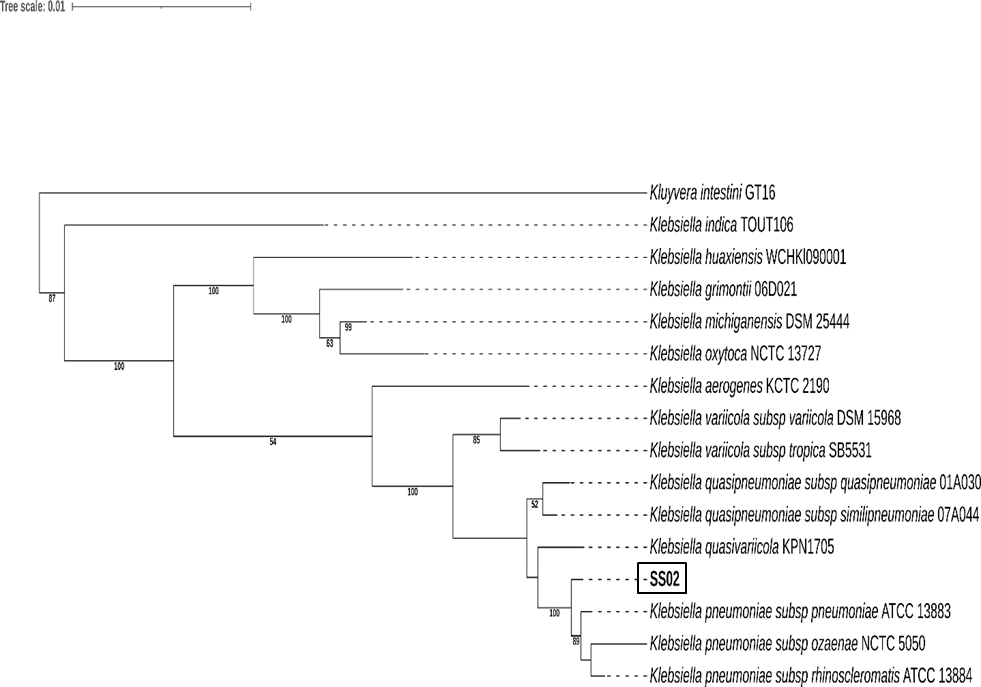

**Fig. S5.** Whole genome-based phylogenetic tree generated in KBase, showing the taxonomic placement of strain SS02 relative to *Klebsiella* type strains. The scale bar indicates substitution per site.

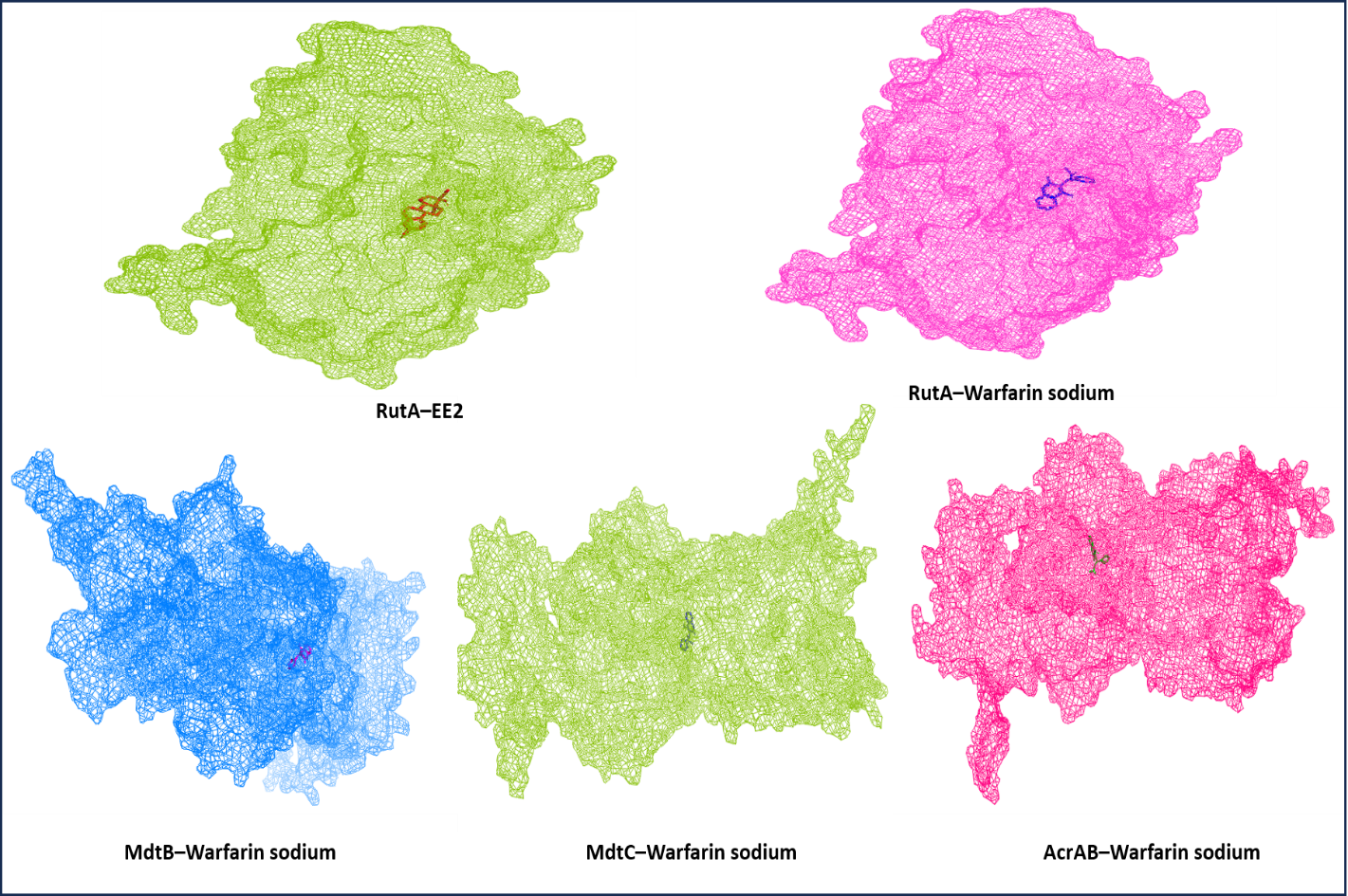

**Fig. S6.** Docked complexes of EE2 and warfarin sodium with predicted protein targets from strain SS02. Surface representations show ligand binding of EE2 and warfarin to RutA (oxidative entry enzymes), and warfarin sodium to RND-type efflux components MdtB, MdtC, and AcrAB systems.

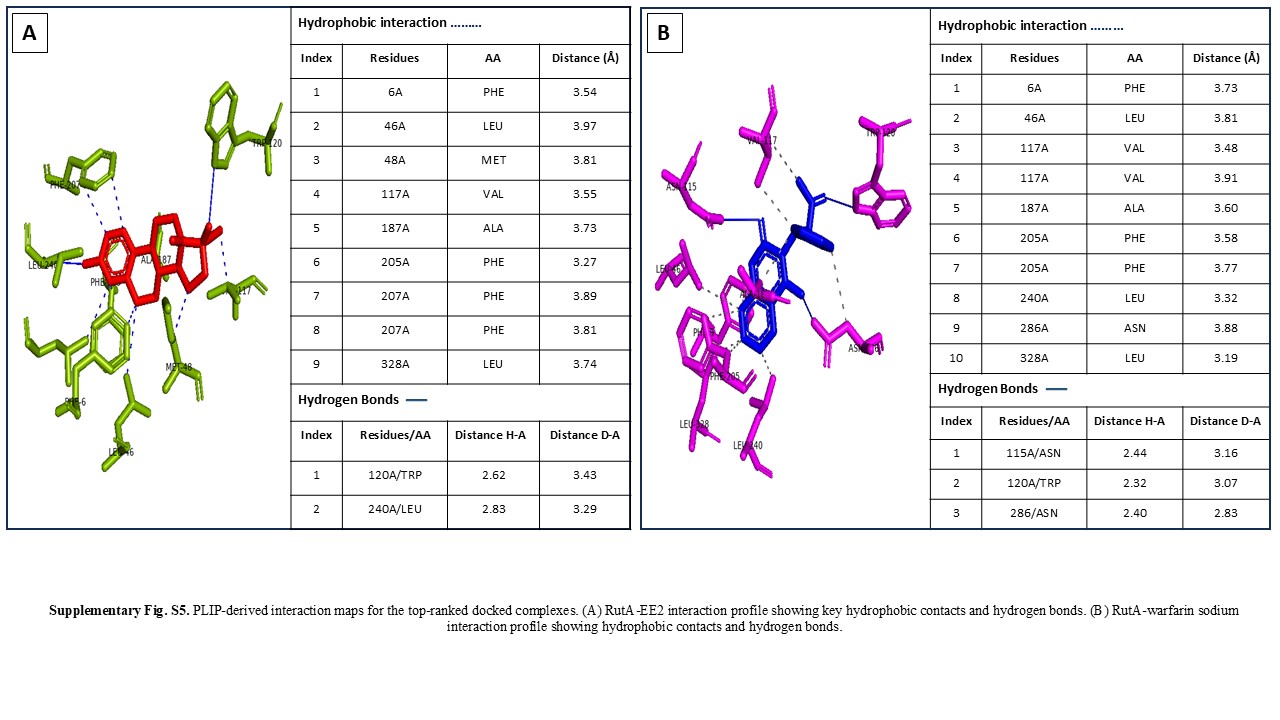

**Fig. S7.** PLIP-derived interaction maps for the top-ranked docked complexes. (A) RutA-EE2 interaction profile showing key hydrophobic contacts and hydrogen bonds. (B) RutA-warfarin sodium interaction profile showing hydrophobic contacts and hydrogen bonds.

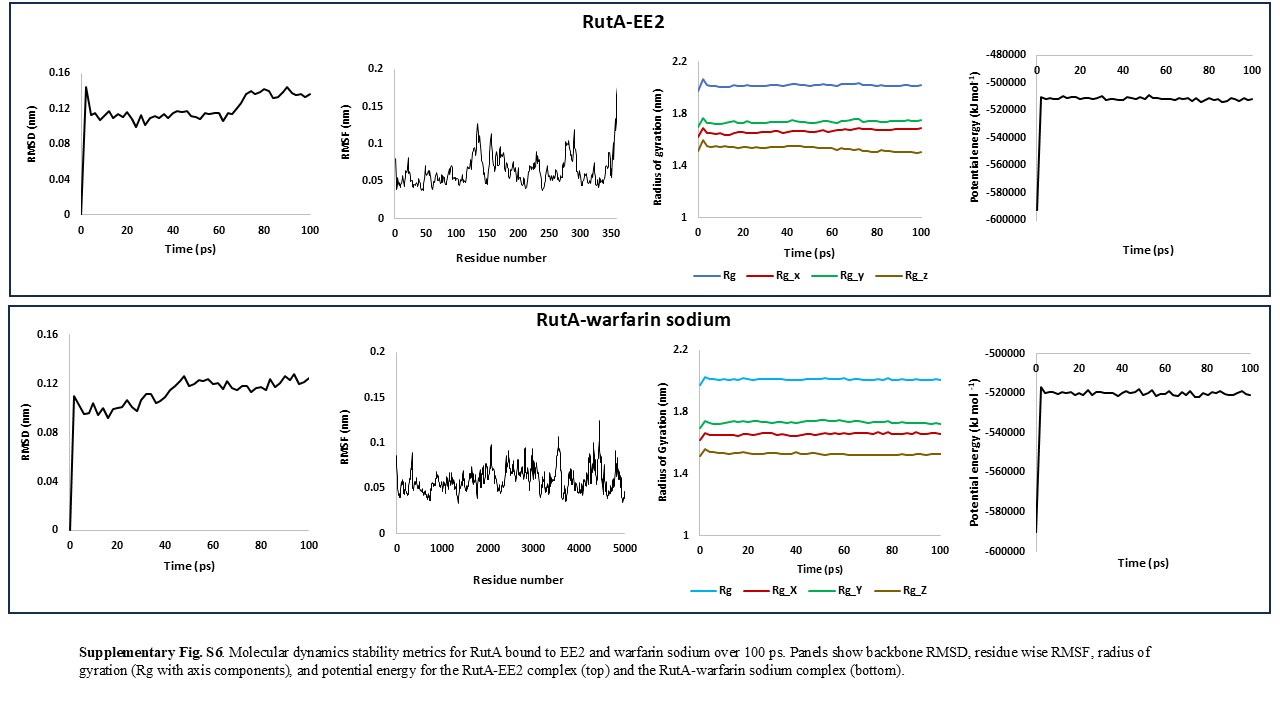

**Fig. S8.** Molecular dynamics stability metrics for RutA bound to EE2 and warfarin sodium over 100 ps. Panels show backbone RMSD, residue wise RMSF, radius of gyration (Rg with axis components), and potential energy for the RutA-EE2 complex (top) and the RutA-warfarin sodium complex (bottom).

**Table S1. Composition of Enrichment media**

| **Composition** | **g L^-1^** |
| --- | --- |
| KH_2_PO_4_ | 3 |
| Na_2_HPO_4_ | 6 |
| NaCl | 5 |
| MgSO_4_ | 0.1 |
| KNO_3_ | 0.875 |
| EE2 | 0.01 |
| Yeast extract | 0.005 |

**Table S2. Composition of Warfarin sodium supplemented MSM agar**

| **Composition** | **g L^-1^** |
| --- | --- |
| KH_2_PO_4_ | 3 |
| Na_2_HPO_4_ | 6 |
| NaCl | 5 |
| MgSO_4_ | 0.1 |
| KNO_3_ | 0.875 |
| Warfarin sodium | 0.02 |
| Agar | 15 |

**Table S3. Composition of growth media**

| **Composition** | **g L^-1^** |
| --- | --- |
| KH_2_PO_4_ | 3 |
| Na_2_HPO_4_ | 6 |
| NaCl | 5 |
| MgSO_4_ | 0.1 |
| KNO_3_ | 0.875 |
| Glucose/ EE2/ Warfarin sodium | 4/ 0.01/ 0.02 |

**Table S4: Summary of the biochemical characteristics and substrate utilization profile of the strain SS02**

| **Biochemical tests** | **SS02** |
| --- | --- |
| Gram staining | Gram negative |
| KOH string test | + |
| Catalase test | + |
| Oxidase test | - |
| Indole test | - |
| Methyl Red | - |
| Voges-Proskauer | + |
| Citrate utilization | + |
| **Substrate utilization tests:** | |
| Lactose | + |
| Xylose | - |
| Maltose | + |
| Fructose | - |
| Dextrose | + |
| Galactose | w+ |
| Raffinose | + |
| Trehalose | + |
| Melibiose | + |
| Sucrose | + |
| L-Arabinose | + |
| Mannose | + |
| Inulin | - |
| Sodium gluconate | w+ |
| Glycerol | + |
| Salicin | + |
| Dulcitol | - |
| Inositol | + |
| Sorbitol | + |
| Mannitol | w+ |
| Adonitol | + |
| Arabitol | + |
| Erythritol | - |
| alpha-Methyl-D-glucoside | - |
| Rhamnose | + |
| Cellobiose | - |
| Melezitose | - |
| alpha-Methyl-D-Mannoside | - |
| Xylitol | - |
| ONPG | + |
| Esculin | + |
| D-Arabinose | - |
| Citrate | + |
| Malonate | + |
| Sorbose | - |

+, Positive; -, Negative; w+, Weakly positive

**Table S5. Candidate genes and functional modules supporting the proposed EE2 and warfarin sodium response pathway in *Klebsiella pneumoniae* SS02.**

| **Gene/protein** | **Annotation** | **Pathway role** | **Linked substrate** | **Evidence source** | **Interpretation** |
| --- | --- | --- | --- | --- | --- |
| RutA | FMN-dependent monooxygenase-like protein / pyrimidine monooxygenase | Shared oxidative entry | EE2 + warfarin sodium | PGAP, RAST, eggNOG, conserved domain analysis, docking, MD simulation | Proposed common entry enzyme for oxidative activation of both substrates |
| RutR | TetR-family transcriptional regulator | Regulatory control of Rut-linked response | EE2 + warfarin sodium | Genome annotation, RutR-operator docking, PLIP interaction analysis | Supports ligand-associated derepression model through weakened DNA binding |
| HpaB/HpaC | 4-hydroxyphenylacetate hydroxylase system / flavin-dependent hydroxylase components | Aromatic hydroxylation and redox-supported oxidation | EE2/warfarin-derived intermediates | PGAP, RAST, eggNOG | Supports downstream oxidative processing of aromatic intermediates |
| Oxidoreductases | NAD(P)H-dependent oxidoreductases | Redox processing | EE2 + warfarin sodium | Genome annotation, STRING network | Supports electron-transfer reactions linked to oxidative metabolism |
| Dehydrogenases | Alcohol/aldehyde/short-chain dehydrogenases | Intermediate oxidation | EE2/warfarin-derived intermediates | PGAP, RAST, eggNOG | Supports conversion of partially oxidized intermediates |
| Hydrolases | Putative esterases/hydrolases | Hydrolytic transformation of intermediates | Warfarin sodium and aromatic intermediates | Genome annotation | Supports cleavage or modification of transformed pharmaceutical intermediates |
| Cat genes | Catechol pathway enzymes | Ring cleavage and aromatic funneling | Aromatic intermediates | RAST, eggNOG | Supports entry of aromatic metabolites into central metabolism |
| Pca genes | Protocatechuate pathway enzymes | Aromatic ring cleavage and β-ketoadipate route | Aromatic intermediates | RAST, eggNOG | Supports downstream conversion into central carbon metabolism |
| Ben genes | Benzoate pathway enzymes | Benzoate/aromatic compound metabolism | Warfarin-related aromatic intermediates | RAST, eggNOG | Supports metabolism of aromatic side-chain products |
| β-ketoadipate pathway genes | β-ketoadipate-associated enzymes | Central aromatic funneling route | EE2/warfarin-derived aromatic intermediates | Genome annotation | Supports conversion of ring-cleavage products into central metabolism |
| mdtABC | RND-type multidrug efflux system | Efflux-assisted tolerance | Warfarin/pharmaceutical stress | Genome annotation, PAβN inhibition assay | Supports survival under elevated warfarin sodium stress |
| acrAB-tolC | RND multidrug efflux system | Multidrug/pharmaceutical efflux | Warfarin/pharmaceutical stress | Genome annotation, PAβN inhibition assay | Supports export-linked defense during pharmaceutical exposure |

**Table S6. PyMOL-based interface contact analysis of apo and ligand-bound RutR–operator DNA complexes.**

| **Model** | **Regulatory state** | **Protein atoms within 4 Å of DNA** | **DNA atoms within 4 Å of protein** | **Residue-level contacts** | **Interpretation** |
| --- | --- | --- | --- | --- | --- |
| Apo RutR-DNA | OFF state | 100 | 102 | 227 protein atoms and 479 DNA atoms | Strong operator binding |
| EE2-bound RutR-DNA | ON-like state | 71 | 76 | 138 protein atoms and 351 DNA atoms | Reduced DNA engagement |
| Warfarin-bound RutR-DNA | ON-like state | 79 | 85 | 153 protein atoms and 351 DNA atoms | Reduced DNA engagement |

**Table S7. In silico mutation of selected RutR DNA-contacting residues and predicted effects on RutR-operator DNA interaction.**

| **Sequential position** | **WT residue** | **Mutated residue** | **Mutation notation** | **Expected structural consequence** |
| --- | --- | --- | --- | --- |
| 34 | Phenylalanine | Leucine | F34L | Loss of aromatic side chain; reduced aromatic/hydrophobic contribution at residue 34. |
| 37 | Histidine | Asparagine | H37N | Loss of cationic or partially cationic phosphate-anchoring potential; possible redistribution of hydrogen bonding through Asn37. |
| 47 | Arginine | Cysteine | R47C | Loss of strongly positive guanidinium group; expected disruption of phosphate-backbone salt-bridge anchoring. |
| 52 | Lysine | Glutamine | K52Q | Loss of positively charged lysine side chain; WT Lys52-associated phosphate salt bridge was absent in the mutant PLIP profile. |
| 113 | Lysine | Glutamic acid | K113E | Charge reversal from positive to negative; WT Lys113-associated phosphate salt bridge was absent in the mutant PLIP profile. |
| 178 | Threonine | Serine | T178S | Conservative polar substitution; removal of one methyl group may slightly alter local packing or hydrogen-bond geometry. |

**Note S1. Biosafety profile of strain SS02:**

The biosafety profile of strain SS02 was evaluated through combined virulence, resistance, and genome defense analyses. Although in silico prediction associated the strain with human pathogenic lineages, this signal is likely driven by phylogenetic proximity within the *Klebsiella pneumoniae* complex rather than the presence of defining pathogenic determinants. No Shiga toxin genes or major hypervirulence regulators were detected. The identified virulence-associated features, including *fimH*, *mrkA*, *fyuA*, *irp2*, *iutA*, *traT*, and *clpK1*, represent widely conserved fitness and stress adaptation traits that support environmental persistence, adhesion, and iron acquisition, rather than direct host invasion. The strain exhibits a multidrug resistance repertoire; however, these traits align with environmental exposure to chemical stressors and are consistent with the observed efflux-mediated tolerance mechanisms linked to xenobiotic survival. Importantly, the genome encodes a functional Type I restriction–modification system along with additional methyltransferases, which provide sequence-specific recognition and restriction of foreign DNA, thereby limiting horizontal gene acquisition and contributing to genomic stability. The absence of Type III and Type IV restriction systems suggests a controlled restriction landscape without extensive methylation-dependent targeting. Taken together, the lack of key toxin systems, the predominance of generalist fitness determinants, and the presence of intrinsic genome defense mechanisms support the classification of SS02 as suitable for controlled laboratory investigations under standard biosafety level 2 practices, while cautioning against uncontained environmental application.
